# The Structural Basis of alpha/beta-tubulin Assembly and Disassembly by Tubulin Cofactors

**DOI:** 10.64898/2025.12.30.697104

**Authors:** Aryan Taheri, Md Ashaduzzaman, Vishv Gill, Jawdat Al-Bassam

## Abstract

Microtubules polymerize from cytoplasmic pools of soluble αβ-tubulin heterodimers that support diverse cellular functions. The tubulin cofactors, TBCC, TBCD, TBCE, and the Arl2 GTPase, form TBC-DEG assemblies that regulate αβ-tubulin assembly and disassembly from α- and β-tubulins, yet their underlying mechanisms remain incompletely understood. Here, we reconstitute the human TBC-DE and TBC-DEG assemblies from eukaryotic cells co-purified with monomeric β-tubulin intermediates and determine their cryo-EM structures. The structures reveal that TBC-DEG disassembles αβ-tubulin by releasing α-tubulin through a lever-arm-like rotation in TBCE coupled to major conformational change in Arl2 upon its nucleotide release, while TBCD tightly holds β-tubulin. TBCD dissociates α-tubulin by refolding the β-tubulin H10-S8 loop at its intradimer interface. The TBC-DEG-β-tubulin or TBC-DE-β-tubulin assemblies undergo extensive back-to-back dimerization mediated by β–β-tubulin homodimers, formed through their dissociated H8 helices at unoccupied intradimer interfaces. Structural comparisons demonstrate that the TBCE mechanical rotation, driven by the Arl2 GTPase cycle, either delivers α-tubulin or removes it from beneath the TBCD-bound β-tubulin and is directionally regulated by TBCC stabilizing αβ-tubulin interfaces. Our findings suggest that TBC-DEG/TBCC catalyzing heterodimerization of α-tubulin with β-tubulin may have evolved to counteract the β-tubulin intrinsic tendency to form off-pathway toxic homodimers through its exposed α-tubulin-binding intradimer interface.

## Introduction

Microtubules (MTs) are dynamic polymers that drive cell division, serve as tracks for intracellular transport, and form the cytoskeletal cores of motile cilia and flagella(*1, 2*). MTs polymerize from soluble αβ-tubulin heterodimers, which are maintained in concentrated pools in the cytoplasm(*1–3*). The cellular concentrations of soluble αβ-tubulin are autoregulated through feedback mechanisms involving the translation regulatory protein, SCAPER, and the tubulin-specific translation corepressor,TTC5, which uniquely co-regulate ribosome translation of tubulins(*4, 5*). Newly synthesized α- and β-tubulin polypeptides are captured by prefoldin assemblies and delivered to inside the chambers of the chaperonins (TRiC/CCT1-8), where they are folded through successive ATP hydrolysis cycles(*6*).

Following chaperonin mediated folding, the tubulin cofactors (TBCs) function as tubulin-specific chaperones that promote the assembly of newly folded α- and β-tubulin monomers into αβ-tubulin heterodimers(*7–9*). This process is mediated by five highly conserved TBC subunits TBCA, TBCB, TBCC, TBCD, TBCE, and the essential Arf-like 2 (Arl2) GTPase (*9, 10*). Biochemical reconstitution of the yeast TBCD, TBCE, Arl2 revealed their shared role as multi-subunit assembly (*9, 10*). TBCD, TBCE, and Arl2 form heterotrimeric TBCD-TBCE-Arl2 assemblies, termed TBC-DEG, which binds to individual αβ-tubulin, associates with TBCC to trigger Arl2 GTP hydrolysis as GTPase activating protein (GAP) (*9, 10*). Arl2 GTP hydrolysis cycles within TBC-DEG are thought to regulate α- and β-tubulin assembly into αβ-tubulin heterodimers (also termed biogenesis) and its disassembly (also termed degradation) (*9, 10*). The critical importance of this pathway is underscored by its involvement in human disease: disruption of αβ-tubulin dimerization causes neurological and developmental disorders, including polymicrogyria, pachygyria, and lissencephaly(*11–13*). Similarly, mutations in TBCD, TBCE, or Arl2 lead to neurological and developmental disorders such as infantile encephalopathy and corpus callosum hypoplasia (*14, 15*), Kenny–Caffey or hyperparathyroidism syndrome, (*16–18*); and microcornea, rod–cone dystrophy, cataract, and posterior staphyloma (MRCS) syndrome(*19*), respectively.

Cryo-EM structures of yeast TBC-DEG in multiple states revealed pre- and post-catalytic conformations bound to αβ-tubulin, reconstituted in vitro(*20*). Recent human TBC-DEG structures uncovered parallel structural states of the human TBC assemblies(*21*). In the pre-catalytic states, TBC-DEG forms a heterotrimeric, cage-like organization that encases a single αβ-tubulin heterodimer, with TBCD encircling β-tubulin while TBCE binds along the α- and β-tubulin(*20*). In post-catalytic states, yeast and human TBC-DEG/TBCC–αβ-tubulin show TBCC engaging TBC-DEG and αβ-tubulin through multi-domain, long-range interactions(*20, 21*). In post-catalytic states, TBC-DEG/TBCC–αβ-tubulin complexes show TBCC C-terminal (TBCC-C) β-helix GAP domain binds to GTP-loaded Arl2 and interacts with the N-terminal region of TBCD(*20*). In the second post-catalytic state, the TBCC linker domain (TBCC-L), which connects its N- and C-terminal domains, binds along TBCD, positioning the TBCC N-terminal three-helix bundle (TBCC-N) beneath TBCD to form a wedge that recognizes and stabilizes the native αβ-tubulin conformation(*20, 21*). Yeast structures also show TBCE disengaging from αβ-tubulin while remaining associated with TBC-DEG and rotating along the longitudinal αβ-tubulin interface, while the human structures suggest TBCE may dissociate fully from TBC-DEG after this step(*20, 21*).

To investigate the catalytic transitions of TBC assemblies bound to α- and β-tubulin intermediates in a physiological context, we have reconstituted human TBCD, TBCE, and Arl2 in eukaryotic cells and purified the resulting assemblies together with de novo tubulin-bound intermediates. Co-expression of TBCD and TBCE, with or without Arl2, yielded TBC-DE and TBC-DEG assemblies that recruited native β-tubulin monomers *de novo*. Cryo-EM structures of TBC-DEG-β-tubulin compared to TBC-DEG-αβ-tubulin reveal the dissociation of α-tubulin is driven by a major lever-arm-like outward rotation of TBCE coupled to large-scale conformational changes in the Arl2 GTPase domain upon nucleotide release. The structures show that TBCD promotes α-tubulin release by refolding the ∫-β-tubulin H10-S8 loop resulting in a disordered H8 helix at its unoccupied intradimer interface. The β-tubulin monomer induces the TBCD C-terminal domain to stabilize the outward-swung TBCE conformation. Both TBC-DE-β-tubulin and TBC-DEG-β-tubulin exhibit a propensity to form back-to-back dimers, mediated by the β-tubulin unoccupied intradimer interfaces to form β–β-tubulin homodimers.

Together with previous structural studies(*20, 21*), we present a complete, multi-step catalytic cycle for αβ-tubulin biogenesis and degradation. In this cycle, TBC-DEG forms a “vise” that mediates αβ-tubulin heterodimer disassembly by TBCD tightly gripping β-tubulin, releasing α-tubulin, while Arl2 GDP dissociation promotes TBCE swinging outward to displace α-tubulin. Subsequent GTP binding to Arl2 promotes the inward swing of TBCE, enabling α-tubulin loading beneath the TBCD bound β-tubulin while TBCE dissociates from αβ-tubulin but not from TBCD. These structural intermediates suggest that TBCC-dependent, GTP-hydrolysis–drives αβ-tubulin assembly pathway on TBC-DEG, which may have evolved to counteract β-tubulin’s intrinsic propensity to form β–β homodimers in the absence of α-tubulin.

## Results

### Human TBC-DEG and TBC-DE assemblies recruit β-tubulin monomers from eukaryotic cells

To reconstitute human tubulin cofactors in near native cellular system with native α- and β-tubulin biogenesis, we co-expressed human TBCD, TBCE, and Arl2, or human TBCD and TBCE, using a baculovirus expression system in *Trichoplusia ni* (Tni) cells (**Figure S1A–C; Figure 1A**). We purified intact TBC assemblies using a StrepII tag on the TBCD N-terminus under near-physiological ionic conditions (pH 7, 150 mM KCl) and in the presence of micromolar GTP, as previously described for yeast TBC-DEG assemblies(*10, 20*) (**Figure S1A**). Size-exclusion chromatography analysis revealed purified assemblies containing stoichiometric human TBCD, TBCE, and Arl2, forming heterotrimeric human TBC-DEG assemblies like those from yeast (**Figure S1A-C; Figure 1A**). These human TBC-DEG assemblies co-purified with a 50-kDa protein identified by mass spectrometry as *Tni* β-tubulin, which is 99% homologous to the human β-tubulin orthologs (**Figure 1A; Figure S1C**). Surprisingly, no α-tubulin was detected co-purifying with these assemblies. When TBCE was omitted from co-expressions, TBCD and Arl2 were insoluble, but in contrast, omitting Arl2 yielded soluble human TBC-DE assemblies that also co-purified with *Tni* β-tubulin. Both human TBC-DEG–β-tubulin and TBC-DE–β-tubulin complexes formed a minor dimeric complex peak as observed by size-exclusion chromatography and mass photometry **(Figure S1B-E)**. Size exclusion chromatography of concentrated human TBC-DEG–β-tubulin analyzed immediately versus 24 hours post-purification revealed the formation of dimeric assembly peak increased over time (**Figure S1F-G**). Mass photometry of fractions representing the monomer, dimer and intermediate peaks confirm assemblies form dimeric TBC-DEG-β-tubulin and similar analyses support this for TBC-DE-β-tubulin (**Figure S1H**). These biochemical co-expression and reconstitution studies show that human TBCD, TBCE, and Arl2 form assemblies orthologous to those in yeast. Unlike yeast TBC-DEG, which is insoluble without Arl2, human TBCD and TBCE co-purify as stable TBC-DE assemblies, indicating that the human system can tolerate the absence of Arl2 but not TBCE.

**Figure 1.**
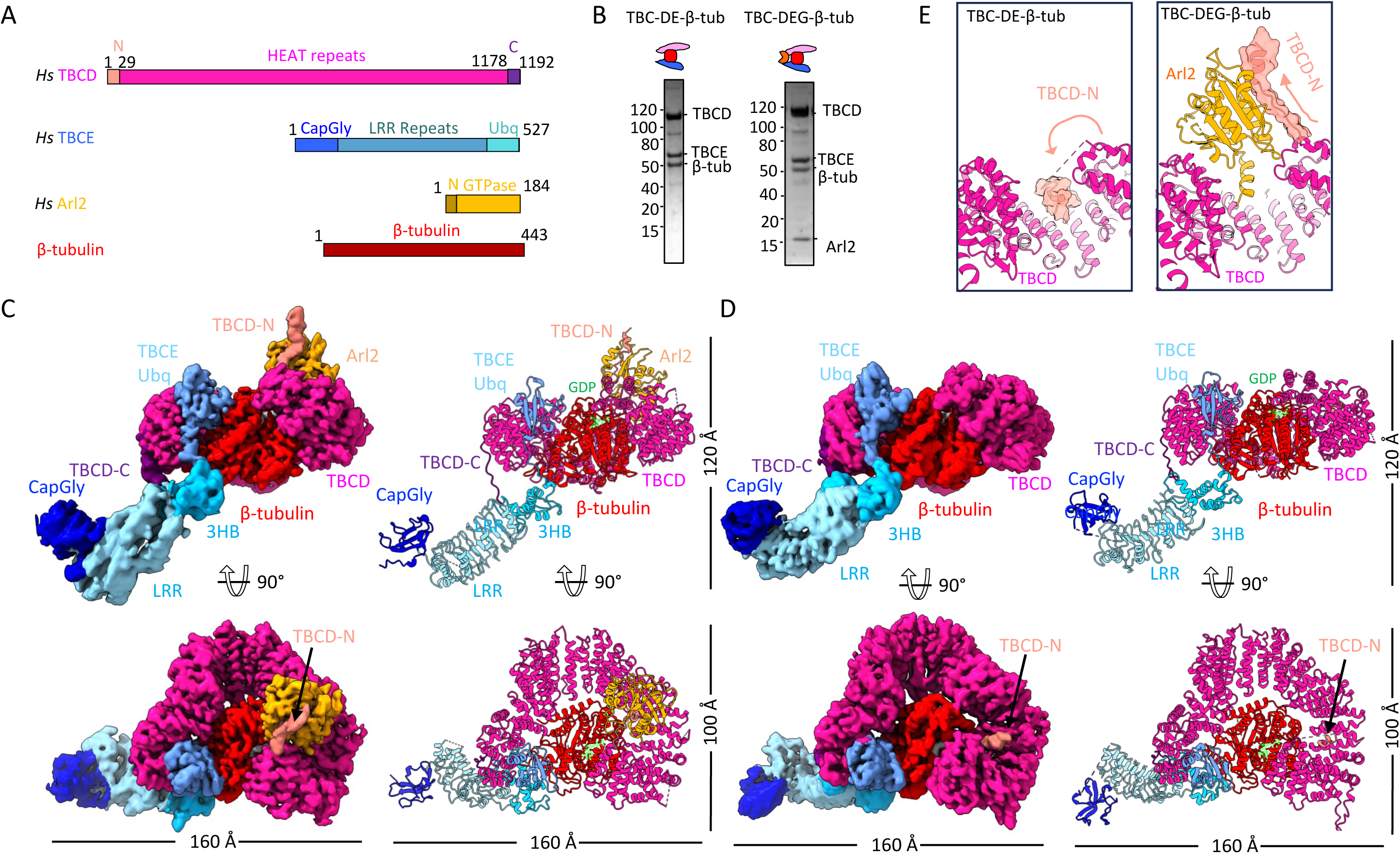
Reconstitution and cryo-EM structures of human TBC-DEG–β-tubulin and TBC-DE–β-tubulin. A) Linear domain organizations of TBC-DEG subunits with domains colored as described in C and D: TBCD (top panel), TBCE (second panel), Arl2 (third panel), β-tubulin (fourth panel). B) Representative SDS–PAGE analysis of purified TBC-DEG–β-tubulin (left) and TBC-DE–β-tubulin (right). See Figure S1 for details. C) *Left panels:* Cryo-EM maps of TBC-DEG–β-tubulin segmented and colored according to the domain diagrams in panel A. The lower panel shows a 90° rotated view. See Figure S2 for reconstruction details. *Right panels:* Corresponding ribbon models shown in equivalent orientations. See Figure S3 for additional information. See Figures S4, S7 for details. D) *Left panels:* Cryo-EM maps of TBC-DE–β-tubulin segmented and colored as in panel A, with a lower 90° rotated view. *Right panels:* Ribbon models shown in matching orientations. See Figures S5 and S7 for details. E) Close-up, side view of the TBCD N-terminal turret. *Left:* Interaction between TBCD-N α0 (salmon) and the TBCD turret (pink) in the TBC-DE–β-tubulin complex. *Right:* Interaction of TBCD-N α0 and the adjacent strand (salmon) with the Arl2 GTPase (orange) bound to the TBCD turret (pink) in the TBC-DEG–β-tubulin complex. See Figure S8 for details.

### Cryo-EM structures of human TBC-DEG-β-tubulin and TBC-DE-β-tubulin

To determine the structural organization of TBC-DEG-β-tubulin and TBC-DE-β-tubulin complexes, we used single particle cryo-EM to resolve their native states (**Figures S2A; Figure S3A)**. We prepared samples of TBC-DEG-β-tubulin immediately after purification (Dataset 1; **Figure S2A**) or concentrated TBC-DEG-β-tubulin after 2 hours of incubation with bi-specific crosslinker (Dataset 2; **Figure S2A**). In contrast, TBC-DE-β-tubulin dataset from concentrated samples without using crosslinking (**Figure S3A**). Consistent with our biochemical analyses (**Figure S1**), the predominant species for TBC-DEG-β-tubulin and TBC-DE-β-tubulin were homogeneous heterotrimeric globular assemblies bound to β-tubulin monomers, featuring a single arm-like extension (**Figures S2A; Figure S3A**). 2D-classification also identified dimeric assemblies, corroborating our biochemical observations (**Figure S2A; Figure S3A**). TBC-DEG-β-tubulin dataset2 showed predominately in back-to-back dimeric arrangements (**Figure S2A**). In contrast TBC-DE-β-tubulin dataset resulted in monomeric globular assemblies with single arm-like extension, and a mixture of side-to-side and back-to-back dimeric assemblies (**Figure S3A**).

3D-classification and focused refinement produced medium- to high-resolution TBC-DEG-αβ-tubulin and TBC-DE-αβ-tubulin maps for their globular cores, and lower resolution density in the peripheral arm-like regions (**Figures S2 and Figure S3**). Refined structures included 3.6 Å for the core of TBC-DEG-β-tubulin (**Table 1; Figure S2B**), 4.3 Å TBC-DEG-β-tubulin full assembly with arm-like extension (**Figure S2C**), 4.8 Å for TBC-DE-β-tubulin with the arm-like extension (**Table 1; Figure S3B**). The dimeric assemblies show preferred particle orientations, yielding anisotropic resolution for these maps, which lacked average conformations for the arm-like extension densities. These maps include: 6.3 Å for the back-to-back TBC-DEG-β-tubulin dimers (**Figure S2D**), 6.5-Å for back-to-back TBC-DE-β-tubulin dimers, and 7.6 Å for side-to-side TBC-DE-β-tubulin dimers (**Figure S3C-D**). These maps enabled atomic modeling of the core regions using AlphaFold-predicted structures of TBCD, TBCE, Arl2, and β-tubulin subunits, guided by homology to yeast TBC-DEG assemblies and bear some similarity to a state of the human TBC-DEG-β-tubulin assemblies (**Figure 1C-D; Figure S4; Figure S5)** (*22*). The arm-like density, attributable to the TBCE region, exhibited lateral flexibility, resulting in 5–7 Å resolution distant from the core (**Figure 1C-D, S6**). Modeling of the back-to-back assemblies of TBC-DEG-β-tubulin and TBC-DE-β-tubulin was facilitated by effectively placing each protomer assembly into their monomer densities within each dimer assembly and modeling the contact densities between them (**Table 1**; **Figure S7**).

**Table 1:**
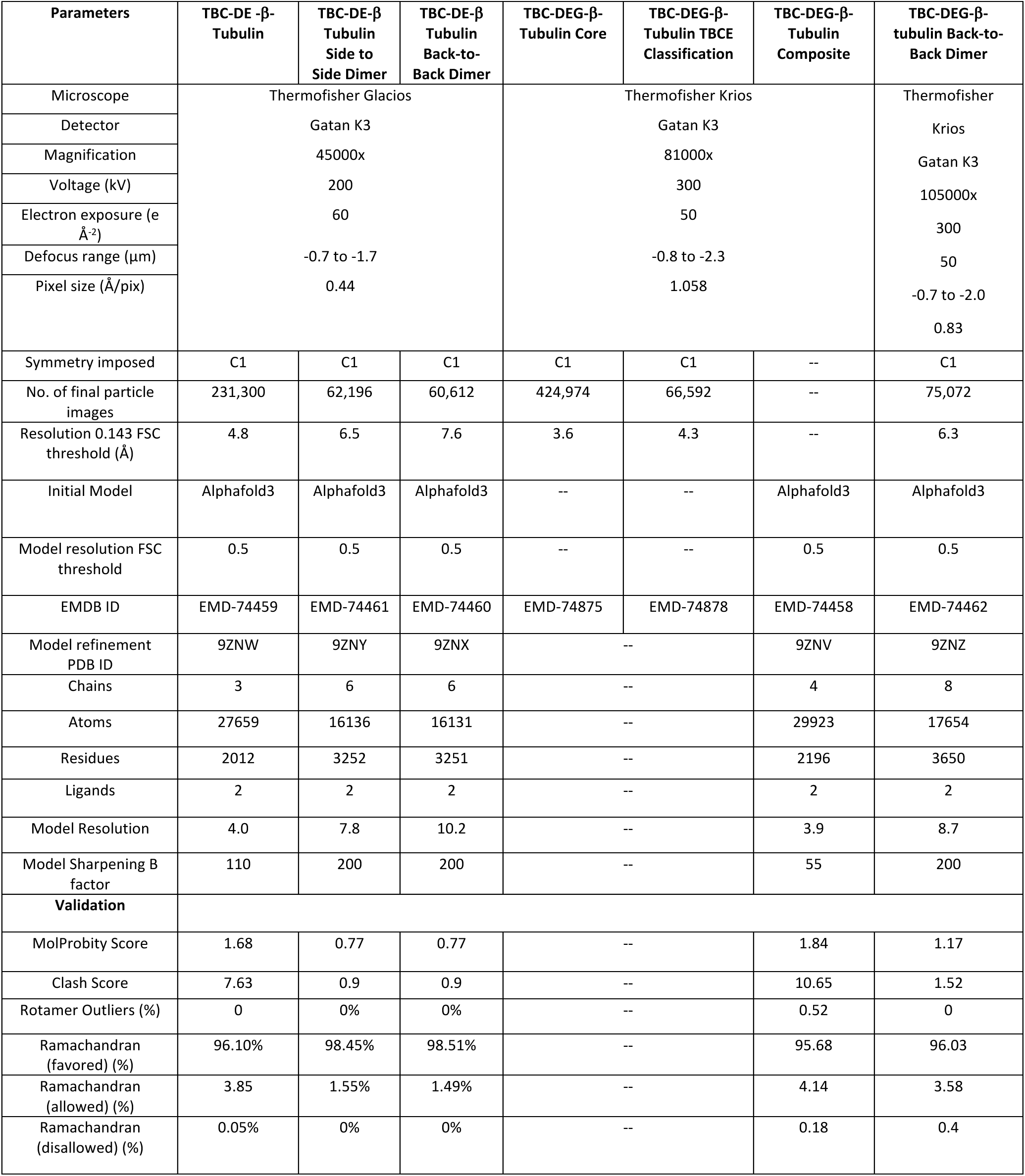
Cryo-EM data collection and model refinement statistics.

### Human TBC-DEG and TBC-DE structures reveal a conserved organization and unique states

The cryo-EM structures of human TBC-DEG–β-tubulin and TBC-DE–β-tubulin define a conserved heterotrimeric architecture of TBCD, TBCE, and Arl2, matching the overall organization observed in the yeast TBC-DEG complex, and explain how the human TBCD and TBCE form stable assemblies in the absence of Arl2 (**Figure 1C-D; Figure S7)**. Human TBCD, although slightly larger than its yeast ortholog, contains 26 α-helical HEAT (<u>H</u>untingtin, <u>E</u>longation factor 3, protein phosphatase 2<u>A</u> and the yeast kinase <u>T</u>OR1) repeat, that assemble into a near-circular N-terminal turret, consisting of repeats 1-13, and a C-terminal spiral domain, consisting of repeats 14-26 **(Figure 1C-D)**. The Arl2 GTPase domain binds atop the TBCD N-terminal turret and encircles the Arl2 N-terminal helix (Nh) **(Figure 1C-D; Figure S8)**. Unlike the yeast structures, the human Arl2 GTPase pocket is unoccupied, indicating it is in a nucleotide-free state **(Figure 1C, S4B)**. The TBCD C-terminal spiral engages the TBCE ubiquitin-like domain (TBCE-Ubq) at an interface nearly identical to that in yeast **(Figure S9A)**. Together, TBCD, Arl2, and TBCE-Ubq form a ring around β-tubulin, establishing extensive hydrophobic and electrostatic contacts that are similar to, but more tightly packed, than those observed in yeast (**Figure 2A-C; Figure S9A)**. TBCE-Ubq also contributes a secondary β-tubulin binding site positioned more closely to β-tubulin than in the yeast TBC-DEG–αβ-tubulin states (**Figure 2D-F**). The β-tubulin C-termini are almost fully folded and bound to TBCD and electron density for them were fully modeled in both TBC-DE and TBC-DEG-β-tubulin structures (**Figure S4F; Figure S5F**)

**Figure 2.**
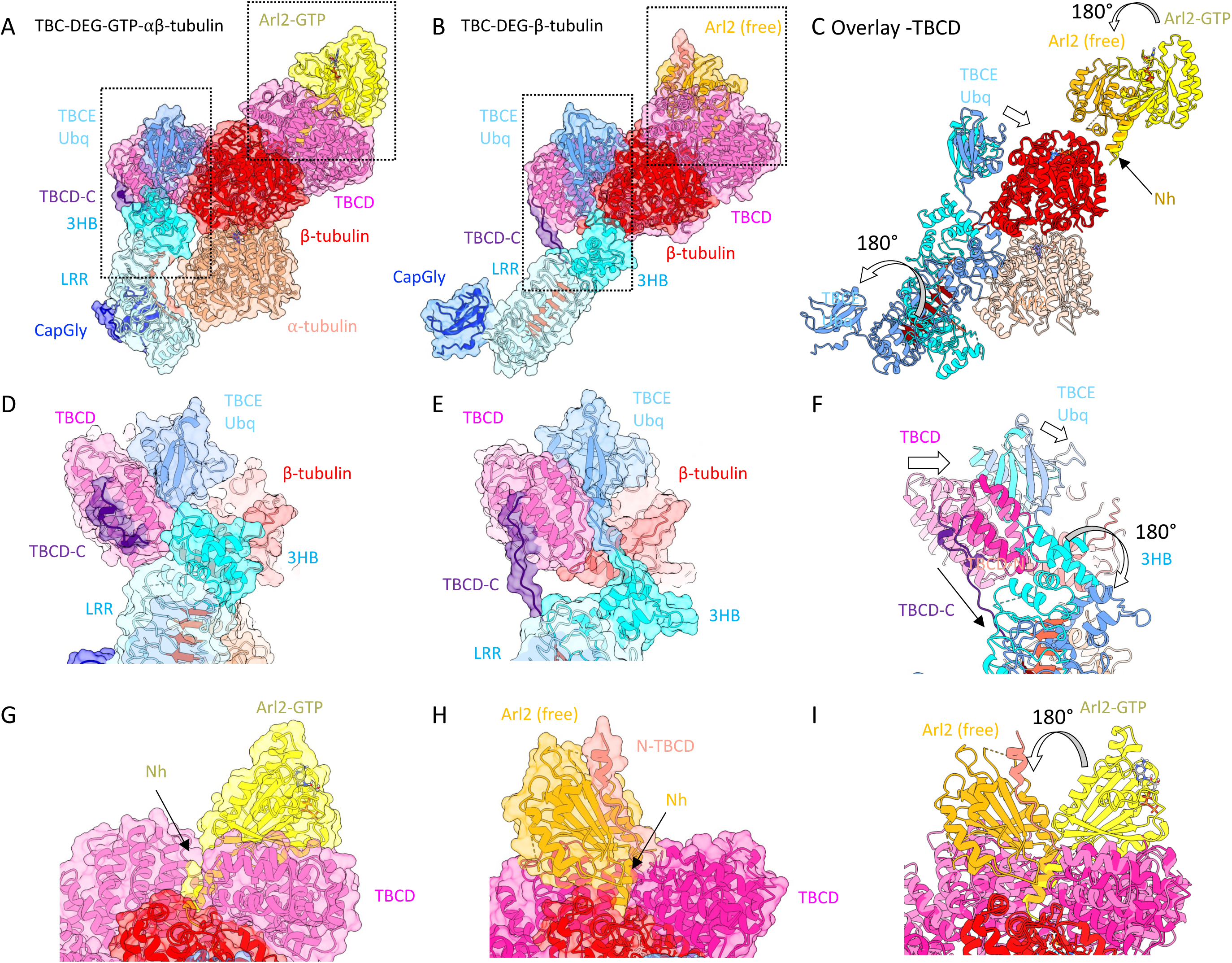
Conformational transitions in TBC-DEG assemblies bound to β-tubulin monomers. A) Side view of yeast TBC-DEG–αβ-tubulin showing TBCD (pink) wrapped around β-tubulin (red) and bound Arl2-GTP (light orange). TBCE contacts both β-tubulin and α-tubulin (salmon). The TBCE-Ubq domain (cyan) and 3HB domain (sky blue) bind β-tubulin, while the TBCE-LRR binds α-tubulin. B) Side view of human TBC-DEG–β-tubulin showing TBCD (dark pink) wrapped around β-tubulin (red) and Arl2 in its nucleotide-free state (orange). The TBCE-Ubq domain (cornflower) and 3HB domain (cyan) contact β-tubulin, with the TBCE-LRR rotated outward. C) Overlay of panels A and B with TBCD removed to highlight Arl2 and TBCE. Arl2-nucleotide-free (orange) and Arl2-GTP (yellow) are shown, together with TBCE from yeast (cyan) and human (cornflower). D) Close-up rotated view of yeast TBC-DEG–αβ-tubulin showing the TBCD C-terminal domain (pink), TBCE-Ubq (cornflower), TBCE-3HB (cyan), and their interactions with TBCD-C (purple) along β-tubulin (red). Arrows indicate rotational movements of TBCE and Arl2. E) Equivalent close-up view of human TBC-DEG–β-tubulin. F) Overlay of panels D and E showing two states of TBCD (light vs. dark pink) and TBCE (cyan vs. cornflower), highlighting conformational transitions. G) Close-up view of the Arl2–TBCD-turret interface illustrating the Arl2-GTP (yellow) H) Close-up view of the Arl2-TBCD-turret illustrating the nucleotide-free Arl2 (orange) I) Overlay of panels G and H showing the ∼180° rotation of the Arl2 GTPase domain (arrow).

The TBC-DE–β-tubulin structure adopts the same overall conformation as TBC-DEG–β-tubulin but lacks Arl2 **(Figure 1C-D)**. In this state, a short acidic helix (termed α0) at the extreme N-terminus of TBCD occupies a site that nearly replaces the Arl2 Nh-helix **(Figure 1E; Figure S8D-F)**. Comparison of the two human structures reveals a critical regulatory role for a 30-residue N-terminal extension of TBCD (TBCD-N), conserved across metazoans, positioned before HEAT repeat 1 **(Figure S8A-C)**. TBCD-N consists of an N-terminal acidic α0 helix followed by a hydrophobic β-strand linker **(Figure 1C-E)**. In the TBC-DEG–β-tubulin structure, α0 binds on the top surface of the Arl2 GTPase domain, while the β-strand linker interacts with the Arl2 central β-sheet **(Figure S8B)**. In the absence of Arl2, as seen in TBC-DE–β-tubulin structure, α0 instead inserts into a highly basic pocket within the TBCD N-terminal turret adjacent to the Arl2 Nh-binding site **(Figure 1E; Figure S8C-F)**. Overall, the TBC-DEG-β-tubulin and TBC-DE-β-tubulin structures exhibit a high degree of conservation with the yeast structures but reveal a mechanism whereby TBCD-N regulates the TBCD N-terminal turret through occupying an empty binding site normally reserved for Arl2.

### Conformational changes in nucleotide free Arl2 GTPase and outward rotation of TBCE

Comparison of the human TBC-DEG-β-tubulin structure with the yeast TBC-DEG-αβ-tubulin structures revealed major conformational changes in TBCE and Arl2 (**Figure 2A–C**). TBCE comprises five domains: an N-terminal CapGly (CapGly) domain, a central leucine-rich repeat (TBCE-LRR) domain, a three-helical bundle (TBCE-3HB), and a conserved linker (TBCE-L) and a C-terminal Ubiquitin domain (TBCE-ubq). In the yeast TBC-DEG–αβ-tubulin structure, the TBCE-LRR–CapGly arm binds longitudinally along αβ-tubulin(*20*). In contrast, in the human TBC-DEG–β-tubulin structure, this arm undergoes a 180° azimuthal twist and outward swing around its anchoring point at the C-terminal end of TBCD (Figure S9A middle and lower panels). In this swung-out state, the α-tubulin–binding site on the TBCE-LRR–CapGly arm faces outward and is displaced ∼50 Å from its position in the yeast structure (**Figure 2A-C**).

The TBCE-LRR–CapGly arm rotation is mediated by conserved elements and interactions with the TBCD C-terminus (**Figure S9B-C**). The rotation pivot is anchored by the TBCE-Ubq, which binds the inner surface of the TBCD C-terminal HEAT 24-26 repeats and connects to the TBCE-LRR–CapGly arm through the conserved TBCE-L linker (**Figure S9C-D**). The TBCE-L linker runs along the inner face of TBCD HEAT26 and hinges the TBCE-3HB domain. The TBCE-3HB rotates ∼180° while maintaining contact with β-tubulin (**Figure 2D–F**). Concomitantly, the TBCD C-terminal HEAT region and TBCE-Ubq domain move inward, forming tighter contacts with β-tubulin (**Figure 2D–F**). The extreme TBCD C-terminus (TBCD-C), extending beyond the HEAT repeats, forms a polyproline type II helix that binds the outer face of TBCD HEAT 26 repeat and forms a “latch” onto the TBCE-LRR domain, stabilizing the swung-out conformation of the TBCE-LRR-CapGly arm. This region contains extensive conservation of the TBCD-C residues at the extreme C-terminus of TBCD; however, due to decreased resolution the molecular interaction stabilizing this TBCD-C latch remains unclear (**Figure 2D-F; Figure S9C-E**). By contrast, in the yeast TBC-DEG–αβ-tubulin structure, the TBCD C-terminus folds back onto its own C-terminal HEAT repeats (**Figure 2E**). Notably, the swung-out TBCE conformation is nearly identical in both the human TBC-DEG–β-tubulin and TBC-DE–β-tubulin in the absence of Arl2 suggesting this complex may form a β-tubulin storage state (**Figure S5E; Figure S6**).

Arl2 undergoes a large rearrangement in the nucleotide-free state in the human TBC-DEG–β-tubulin structure relative to the yeast TBC-DEG-αβ-tubulin structure. The Arl2 GTPase domain in the nucleotide free state is rotated by approximately 180 degrees relative to its GTP-bound conformation in yeast **(Figure 2G–I; Figure S10A–C)**. The N-terminal Arl2 Nh helix remains anchored in the same configuration in the TBCD N-terminal turret in both states (**Figure 2G; Figure S10A-C**). The rotated Arl2 GTPase domain forms a new set of interactions with the TBCD N-terminal turret, distinct from those in the yeast TBC-DEG-αβ-tubulin complex (**Figure 2I; Figure S10A-C**).

Alignment of the Arl2 subunits by their GTPase domains reveals a major rotation in the Nh to GTPase junction (**Figure S10D**). The GTP-binding loops of Arl2 are disengaged in the nucleotide free state compared to the Arl2-GTP state (**Figure S10D**). In the Arl2-nucleotide free state, a loop facing the lower side of the Arl2 GTPase towards TBCD unfolds to directly to contact TBCD and β-tubulin near the GDP E-site (**Figure S10D**). This interaction is mediated by a conserved human Arl2 Arg 114 which binds β-tubulin Asn 99 (**Figure S10F**). In the Arl2-GTP bound state, the equivalent loop is packed and folded up, where the equivalent yeast Arl2 Arg 119 is bound into an acidic pocket while in the re-oriented conformation of the Ar2l-GTPase-TBCD interface (**Figure S10E-F**). These structures suggest that nucleotide dissociation from Arl2 triggers GTPase-domain reorientation likely promoting the αβ-tubulin disassembly mediated by TBCD and TBCE (**Figure S10G-H**). The similarity of the TBC-DE-β-tubulin structure, which lacks Arl2, to the TBC-DEG-β-tubulin structure suggests that the Arl2-nucleotide free state is likely a low energy state of the TBC-DEG and that Arl2-GTP state may be a higher energy state. This Arl2 GTPase conformational transitions likely rearranges TBCD and the associated TBCE at its C-terminal domain (see below).

### TBCD dissociates α-tubulin from β-tubulin by refolding the H10-S8 loop and disordering the H8 helix

Comparison of the TBC-DEG-β-tubulin structure with the yeast TBC-DEG-αβ-tubulin structure reveals how TBCD dissociates the α-tubulin by reorganizing β-tubulin loops at the intradimer interface (**Figure 3A-F**). In the TBC-DEG-β-tubulin structure, TBCD, TBCE-Ubq more tightly bind to β-tubulin compared to their conformation in the yeast TBC-DEG-αβ-tubulin structure burying almost twice the surface area (4500 Å^2^ versus 2280 Å^2^; **Figure S12**). In the TBC-DEG-β-tubulin and TBC-DE-β-tubulin structures, the C-terminal region of TBCD, comprising HEAT24,25 and 26 repeats, moves inward to engage β-tubulin and refolds its H10-S8 loop (**Figure 3A-C**). Conserved Hydrophobic residues β-tubulin Trp344, Tyr340, and Phe341 in the H10-S8 loop in β-tubulin become captured by TBCD Leu1123, Leu1089 and Phe1123 residues in its inward facing loops and helices (**Figure 3G-I; Figure S8C-D; Figure S11A).** In the native αβ-tubulin and yeast TBC-DEG-αβ-tubulin structure, these same β-tubulin residues in the H10-S8 loop bind hydrophobic residues in α-tubulin stabilizing a major section of the intradimer interface (**Figure 3D-I**). The identical H10-S8 loop refolding observed in the TBC-DE-β-tubulin structure indicates that this conformation represents a low-energy state either in the absence of Arl2 or in its nucleotide-free state.

**Figure 3.**
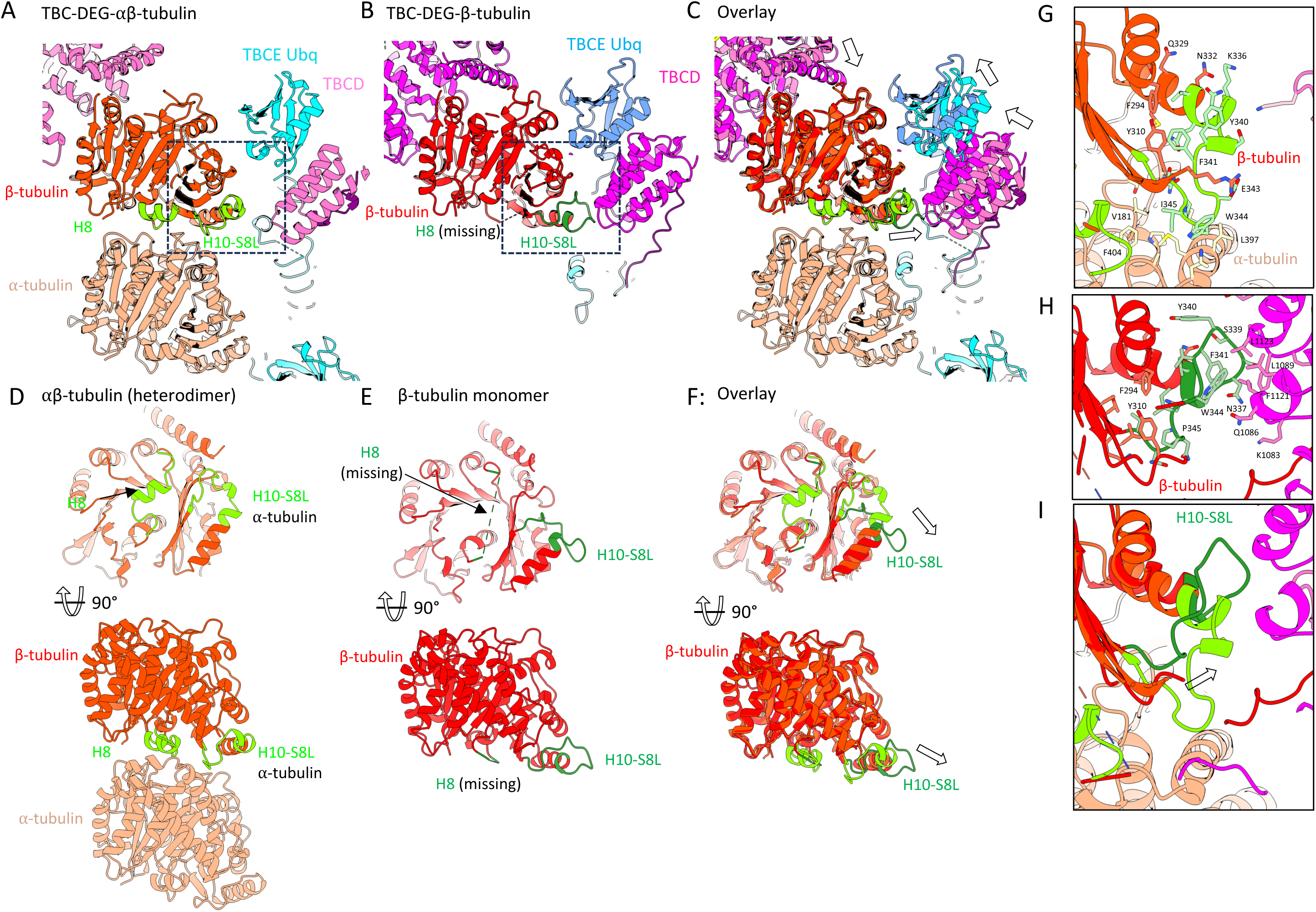
Structural transitions in TBCD that promote β-tubulin dissociation from α-tubulin. A) Cross-section of yeast TBC-DEG–αβ-tubulin. TBCD (pink) wraps around β-tubulin (red) while α-tubulin (salmon) occupies the intradimer interface, marked by the H10–S8 loop and H8 helix (light green). See B) Cross-section of yeast TBC-DEG–β-tubulin showing monomeric β-tubulin (red) with its unoccupied intradimer interface. The H10–S8 loop (dark green) is refolded, and H8 is absent (dotted green line). C) Overlay of panels A and B emphasizing conformational changes in the β-tubulin H10–S8 loop (arrow). D) Two orthogonal views of β-tubulin in the αβ-tubulin. E) Two orthogonal views of β monomer-bound states showing differences in the H10–S8 loop and H8 helix. F) Two orthogonal views of overlays of panels D and E highlighting the structural rearrangements. G) Close-up view of residue interactions within the H10–S8 loop in binding α-tubulin bound αβ-tubulin. H) Close-up view of the residue interaction within H10-S8 loop in monomeric β-tubulin binding TBCD. See Figure S11A for details I) Overlay of panels G and H showing the H10–S8 loop shift (side chains omitted).

The β-tubulin H8 α-helix and its two connecting loops, which directly engage α-tubulin at the N-site GTP in native heterodimers, are completely absent from both TBC-DE-β-tubulin and TBC-DEG-β-tubulin structures (**Figures 3D-F; Figure S11B-C**). This disorder presumably results from loss of α-tubulin contacts. Consistently, reanalysis of recently published human TBC-β-tubulin monomer structures (*21*) reveals that the H8 helix and connecting loops are also absent from the experimental maps, despite being modeled by the authors (**Figures S11C-E**). Together, refolding of the hydrophobic H10–S8 loop combined with H8 helix disorder renders the β-tubulin intradimer interface incompatible with α-tubulin binding.

The conformational changes in TBC-DEG-β-tubulin structure, including the outward rotation of the TBCE LRR-CapGly arm and the Arl2 GTPase domain reorientation, are directly coupled to reorganization of the TBCD C-terminal region, which drives β-tubulin H10-S8 loop refolding (**Figures S9E; Figure S11A**). Binding of the β-tubulin H10-S8 loop to the inner face of HEAT 25-26 repeats induces restructuring of the outward-facing TBCD-C residues, stabilizing TBCD-C elements and latching the TBCE LRR-CapGly arm in its outward-swung position (**Figures S9E; Figure S11A**). These observations indicate that α-tubulin dissociation is mechanistically coupled to the outward rotation of the TBCE LRR-CapGly arm through TBCD C-terminal reorganization.

### The unoccupied β-tubulin intradimer interface promotes β-β-tubulin homodimerization

A substantial proportion of the TBC-DEG-β-tubulin and TBC-DE-β-tubulin particles formed back-to-back dimers, representing ∼10% of assemblies, with this proportion increasing upon extended incubation of the purified complexes, or upon incubation with crosslinker (**Figure S2**). 2D-class averages revealed that TBC-DE-β-tubulin assemblies contain both side-to-side and back-to-back dimers, whereas TBC-DEG assemblies exhibit exclusively back-to-back dimers (**Figures 4A, S3**). We obtained low-resolution structures (6.3-7.2 Å) for the back-to-back dimers of both TBC-DEG-β-tubulin and TBC-DE-β-tubulin, limited by small particle number and strong preferred orientation effects (**Figure S6**). Docking of models derived from higher-resolution TBC-DE-β-tubulin and TBC-DEG-β-tubulin structures enabled interpretation of their organization and assembly interfaces (**Figures 4B-C, S6**).

**Figure 4.**
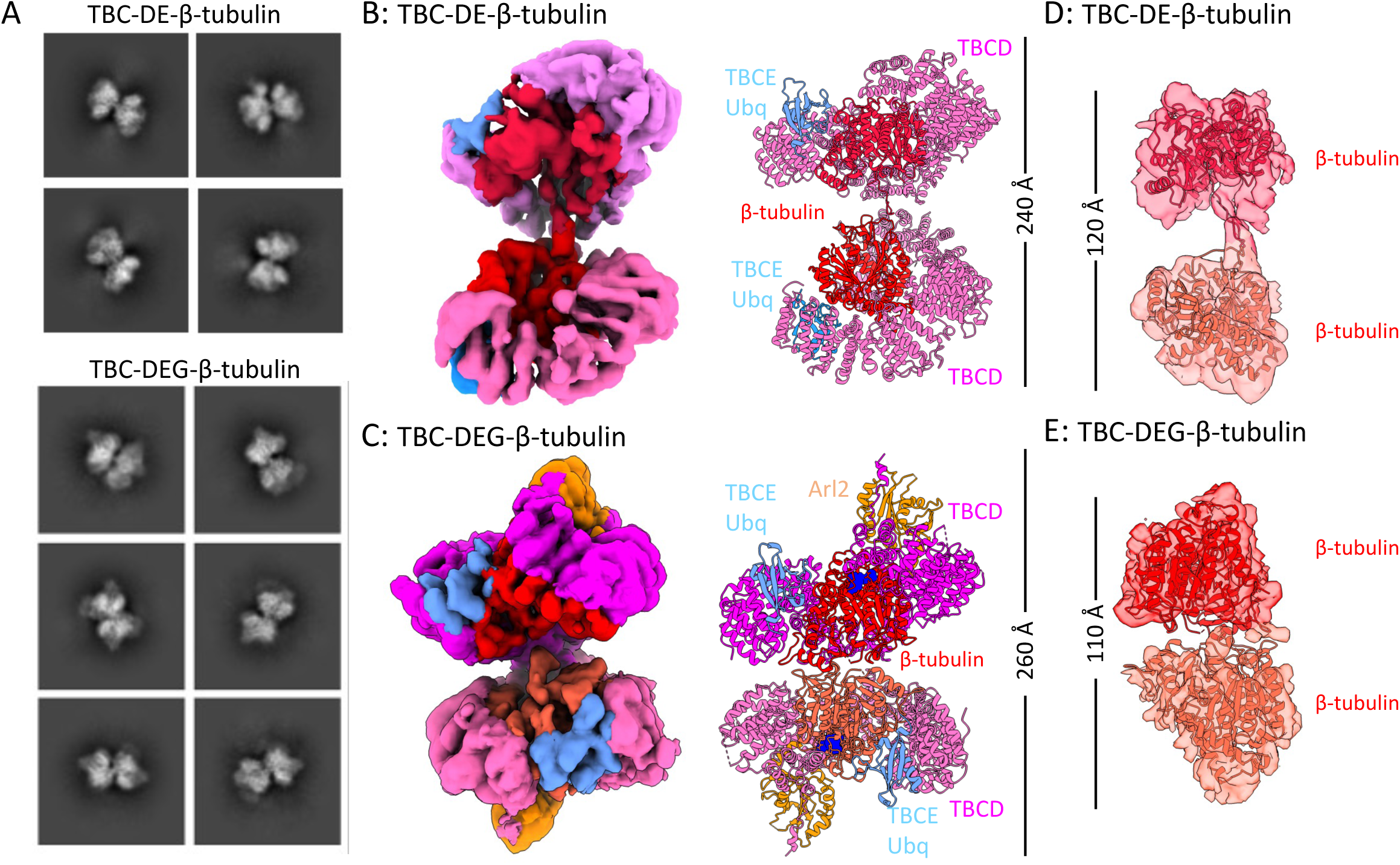
Structural organization of back-to-back TBC assemblies mediated by β–β-tubulin homodimerization. **A)** Representative 2D class averages of back-to-back TBC-DE–β-tubulin dimers (top; see Figure S3) and TBC-DEG–β-tubulin dimers (bottom; see Figure S2). **B)** *Left:* Segmented 6.2 Å reconstruction of the TBC-DE–β-tubulin back-to-back dimer. TBCD (pink) wraps β-tubulin (red), and TBCE-Ubq (blue) binds the TBCD C-terminus. *Right:* Model fitted into the segmented map. See Figure S6 for details. **C)** *Left:* Segmented 7.2 Å reconstruction of the TBC-DEG–β-tubulin back-to-back dimer, with Arl2 (orange) bound to the TBCD N-terminus. *Right:* Corresponding fitted model. See Figure S6. **D)** Segmented densities views of β–β-tubulin homodimers within TBC-DE-β-tubulin back-to-back dimer assemblies. See Figure S6 for more details **E)** Segmented densities views of β-β-tubulin homodimers within TBC-DEG-β-tubulin back-to-back dimer assemblies. See Figure S6 for more details

In the back-to-back dimer configuration, β-tubulin monomers directly participate in dimerization of these assemblies, revealing an intrinsic propensity for β-tubulin to homodimerize via its unoccupied intradimer interface, the same interface normally occupied by α-tubulin in native heterodimers (**Figures 4D-E, S6C-D**). Continuous density extends likely from the H8 helix of one β-tubulin to that of the β-tubulin of the neighboring protomer (**Figure S6**). In contrast, TBC-DE-β-tubulin side-to-side dimers are mediated exclusively by TBCD, which can drive assembly into dimers or higher-order linear oligomers. The combination of these structures and biochemical studies demonstrates that in the absence of α-tubulin, both TBC-DEC-β-tubulin and TBC-DE-β-tubulin undergo spontaneous dimerization, likely mediated by TBCD contacts.

### Mapping the human TBC-DEG disease mutations on the TBC-DEG structure

We mapped pathogenic missense mutations in TBCD, TBCE, and Arl2 from the ClinVar database onto our structural models(*23*) (**Figure S13**). These mutations are distributed throughout the structural regions of TBCD, TBCE, and Arl2, providing insights into disease mechanisms (**Figure S13**). Twenty-seven pathogenic missense mutations are linked to neurodegenerative encephalopathies map across the TBCD structure (**Figures S13A, C**). These mutations predominantly reside within helical regions that mediate hydrophobic packing or at helix termini critical for stabilizing intra- and inter-HEAT repeat interactions, which are essential for β-tubulin binding and encirclement (**Figures S13B-C**). The two TBCE missense mutations associated with hypoparathyroidism, retardation, and dysmorphism (HRD) also termed Kenny-Caffey syndrome localize to the TBCE CAP-Gly domain, likely disrupting its ability to bind the α-tubulin C-terminal tail (**Figure S13B**). In contrast, a mutation causing Giant Axonal Neuropathy resides at the extreme C-terminus of TBCE and likely destabilizes the folding of its TBCE-Ubq domain, impairing its interaction with TBCD. The single Arl2 missense mutation causing microcornea, rod–cone dystrophy, cataract, and posterior staphyloma (MRCS) syndrome localizes to the junction between the Arl2 Nh and GTPase domain, likely disrupting the nucleotide-dependent conformational changes we describe above (**Figures 13B-C**). Our structural mapping reveals how these mutations may impair TBC-DEG assembly, TBCC binding, or conformational transitions in Arl2 which are critical αβ-tubulin biogenesis and degradation. These molecular defects provide a plausible basis for understanding these neurological and developmental disorders that result from disrupted TBC-mediated tubulin homeostasis.

## Discussion

The pathways that maintain soluble, assembly-competent αβ-tubulin are essential for microtubule homeostasis, yet the mechanisms by which tubulin cofactors stabilize monomeric α- and β-tubulin and drive their incorporation into heterodimers have remained incompletely defined. Here, through reconstitution and cryo-EM structural of human TBC-DE and TBC-DEG (Figure 1), we identify previously uncharacterized β-tubulin–bound intermediates that reveal how these assemblies remodel, stabilize, and direct β-tubulin through αβ-tubulin biogenesis and degradation. Both TBC-DEG and TBC-DE capture β-tubulin monomers and induce refolding of the H10–S8 loop leading to the disorder of H8 helix at its unbound intradimer interface (Figures 2–3), generating a state incompatible with α-tubulin binding. The strong structural similarity between TBC-DEG–β-tubulin and TBC-DE–β-tubulin supports β-tubulin recognition as a conserved, obligatory step in the cofactor-mediated αβ-tubulin biogenesis and degradation pathway. TBCD acts as the primary β-tubulin remodeling factor, while TBCE and Arl2 remain constitutively associated, forming a stable triad that functions as an integrated assembly platform. These β-tubulin–bound intermediates protect exposed β-tubulin surfaces that would otherwise promote aberrant self-association. These structures explain how these TBC-DEG and TBC-DE assemblies promote αβ-tubulin disassembly and stabilize β-tubulin monomers.

### A complete model for the TBC-DEG and TBCC mediated αβ-tubulin assembly and disassembly

We propose a comprehensive mechanistic model for the catalytic cycle of TBC-DEG, TBCC, and tubulins during αβ-tubulin biogenesis and degradation (**Figure 5**). This model integrates the yeast and human TBC-DEG-αβ-tubulin and TBC-DEG/TBCC–αβ-tubulin (20)(21), and the β-tubulin intermediates described here, yielding a complete catalytic trajectory (**Figure 5**). TBCD provides the central scaffold, TBCE rotates bidirectionally to position or release α-tubulin, and Arl2 GTP binding/hydrolysis drives these conformational transitions (10, 20). We also identify a TBC-DE–β-tubulin state resembling the nucleotide-free Arl2 conformation of TBC-DEG–β-tubulin, suggesting it serves as a β-tubulin storage or a pre-assembly state (Figure 5; state 0). During the cycle, the nucleotide-free TBC-DEG assembly (Figure 5; State I) loads β-tubulin following its folding (Figure 5; State II). The α-tubulin is also recruited following folding to TBCE through its CapGly domain, which specifically binds its unique C-terminal tail (24) (Figure 5; State III) and then binds the side of α-tubulin via its LLR domain (*21*). GTP binding to the Arl2-GTPase triggers the inward rotation of TBCE LRR-CapGly arm (Figure 5; State IV), positioning α-tubulin beneath β-tubulin, leading to an open αβ-tubulin heterodimer (Figure 5; State V). Subsequent closure of the αβ-tubulin intradimer interface through the rotation of the TBCE LRR-CapGly arm (Figure 5; States VI–VII). TBCC binds to Arl2 GTP via its TBCC-C domain to promote GTP hydrolysis (20). Stabilization of the nascent αβ-tubulin heterodimer is mediated by the TBCC-L and TBCC-N beneath TBCD (Figure 5; State IX) followed by the release of mature αβ-tubulin.

**Figure 5.**
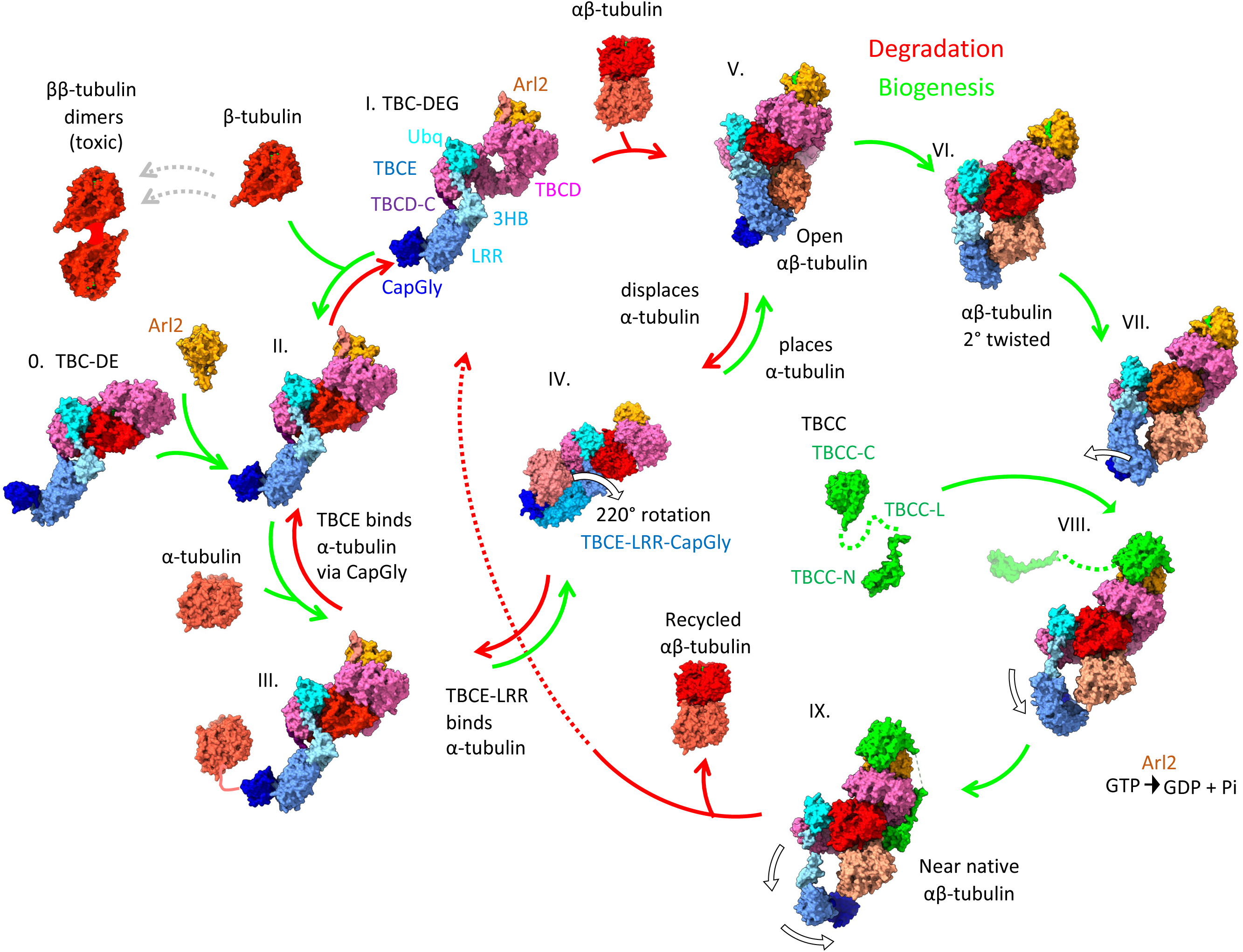
Complete structural model for TBC-DEG/TBCC-mediated αβ-tubulin assembly and disassembly. 0. TBCD and TBCE form TBC-DE complexes that stabilize β-tubulin. I. TBCD, TBCE, and Arl2 assemble into TBC-DEG. II. TBC-DEG binds β-tubulin after folding by CCT or during degradation. Failure to bind α-tubulin rapidly may lead to β–β dimer formation (left dotted arrows). III. TBC-DEG–β-tubulin recruits α-tubulin via the TBCE-CapGly domain. IV. α-tubulin bound along the TBCE-LRR is rotated beneath β-tubulin V. An open αβ-tubulin intradimer interface forms. VI. The interface partially closes into a twisted 2° intermediate. VII. TBCE-LRR–CapGly begins to disengage from α-tubulin. VIII. TBCC binds via TBCC-C to Arl2-GTP. IX. TBCC-C and TBCC-N stabilize the αβ-tubulin interface, and TBCE fully dissociates from α-tubulin. This state appears in both yeast and the reinterpreted human structures (**Figure S14**). Following this step αβ-tubulin and TBCC then dissociate, completing the cycle.

In yeast, TBCC-N binding triggers selective detachment of the TBCE LRR-CapGly arm from αβ-tubulin, while the TBCE-Ubq domain retains stably anchored within the TBC-DEG core (**Figure 5; VIII**)(*20*). In contrast, a recent human TBC-DEG-TBCC-αβ-tubulin structure was interpreted as showing complete TBCE dissociation from TBCD (*21*). However, re-analysis of the deposited human TBCD-Arl2-TBCC-αβ-tubulin cryo-EM maps reveals a prominent, high-occupancy, unmodeled density attributable to the TBCE-Ubq domain based on its fold and location (**Figure S14**). This density is clearly discernible, indicating that TBCE did not dissociate in this conformation (*21*). Rigid-body placement of the TBCE-Ubq domain confirms its presence within the assembly, demonstrating that, consistent with the yeast structures, the TBCE LRR-CapGly arm disengages from αβ-tubulin while the TBCE-Ubq domain maintains its integration into the TBC-DEG core during late-stage αβ-tubulin heterodimer maturation (**Figure 5; IX.**)(*10, 20*).

The structural states reveal that movements of the TBCE LRR-CapGly arm like a mechanical lever arm, potentially analogous to the power stroke of actin-based myosin motor proteins, though with distinct rotational orientation and magnitude(*24*). We show that the pivot point of the TBCE LRR-CapGly arm is the TBCE-3HB attachment to the C-terminal face of TBCD (**Figure 2**; **Figure 5; States IV-V**). GTP binding to Arl2 provides the energy for the inward power stroke, with conformational changes transmitted through the Arl2-GTPase domain to the TBCD scaffold.

### TBC-DEG and TBCC catalysis may have evolved to prevent ββ-tubulin homodimers

A key mechanistic implication of our structures is that monomeric β-tubulin forms β–β homodimer assembly, when its intradimer interface is unoccupied. TBC-DEG–β-tubulin and TBC-DE–β-tubulin both form back-to-back dimers *in vitro* (**Figures 4; Figure S6**), a process enhanced over time (**Figure S1F–H**). The refolded H10–S8 loop, disordered H8 helix, and associated loops contribute to this homodimerization propensity, consistent with features observed in prior β-tubulin monomer structures (21). The disorder of H8 helix likely leads to its swapped association potentially providing an explanation for this dimerization. β-tubulin overexpression in vivo similarly produces toxic oligomers (25), emphasizing the need for cellular mechanisms that suppress this off-pathway reaction. Our data support the idea the TBC-DEG/TBCC machinery may have evolved to transition β-tubulin to assemble with α-tubulin catalytically to prevent the β-tubulin off-pathway intermediates that result from the abundance of β-tubulin monomer intermediates. TBCD stabilizes β-tubulin in a remodeled conformation, while rapid TBCE lever-arm rotation driven by Arl2-GTP positions α-tubulin onto the exposed β-tubulin interface before β–β homodimers can form (Figure 5, left vs. right). This GTP-dependent catalytic cycle likely maintains the fidelity and uniformity of αβ-tubulin heterodimers required for microtubule function.

Together with previous studies (*9, 10, 20, 21*), these findings redefine the tubulin cofactors to catalyze αβ-tubulin biogenesis through a scaffold-based, GTP-powered assembly mechanism. Although many questions remain regarding cellular localization, regulation, and integration with proteostasis pathways, the present work provides a structural and mechanistic foundation for understanding how TBC-DEG and TBCC preserve and maintain concentrated αβ-tubulin pool in the cytoplasm for the homeostasis of MT polymerization across eukaryotes.

## Materials and Methods

### Protein Expression and Purification

Untagged cDNA for Human TBCE (uniprot accession: A0A2R8Y809), Human Arl2 (uniprot accession: P36404) were cloned using Gibson method into Fastbac1 plasmid vector. The untagged into Human TBCD (uniprot accession: Q9BTW9) was cloned in frame with N-terminal StrepII tag into Fastbac1 plasmid vector and verified with sequencing. For baculovirus expression, The Fastbac-strp-TBCD, Fastbac-TBCE, and Fastbac-Arl2 plasmids transformed into DH10bac cells and plated onto LB-agar plates containing bluo-gal and incubated for 72 hours. After 72 hours, white colonies were selected for growth and purification of bacmid for protein expression through isopropanol precipitation. SF9 cells (Expression systems, Davis CA) were transfected at a cell density of 10^6^ cells*mL^-1^ using Cellfectin (Thermofisher) to produce P1 virus with individual Bacmids for Strp-TBCD, TBCE and Arl2. Baculoviruses were collected after five days, and 50mL of 10^6^ cells/mL^-1^ SF9 cells (Expression systems, Davis, CA) were infected to produce P2. After five days, virus was collected, and 50mL of 10^6^ cell/mL ^SF^9 were infected to produce P3. 1L of 10^6^ cell/mL ^cell^ density of Tni cells (Expression systems, Davis, CA) were infected to express protein. Expression of TBC-DEG or TBC-DE was accomplished through infection of the 1L 10^6^ cells*mL^-1^ Hi5 with a ratio of 9 mL TBCD-strep, 3 mL TBCE, and 1 mL of Arl2 P3 virus in the case of TBC-DEG purification. In the case of TBC-DE purification, infection was performed with 9 mL TBCD-strep and 3 mL TBCE. Hi5 cells were incubated for 60 hours, and the cells were pelleted and flash frozen. For protein purification, cells were resuspended in 50mL of lysis buffer (50 mM HEPES, 150 mM KCl, 1 mM MgCl2, 3 mM β-mercaptoethanol, 5% glycerol, with 2 protease inhibitors tablets (Sigma Aldrich). Cell lysates were clarified by centrifugation at 18,000 rpm for 30 min at 4°C. Protein was purified from lysate using 2mL of Streptactin XT beads (IBA-life sciences). Protein was eluted through 5 column volumes of lysis buffer containing 50 mM Biotin. Protein was then concentrated and loaded onto a Superdex-200 10/300 size exclusion column. Fractions of eluted protein were combined, concentrated, aliquoted, and frozen in liquid nitrogen after supplementing each aliquot with an additional 5% glycerol to analyze the biochemical propensity for TBC-DEG-β-tubulin to form dimers, freshly purified TBC-DEG-β-tubulin was concentrated to 2.0 mg/ml then they were either injected immediately or incubated on ice for 24 hours then injected on a Superdex 200 10/300, equilibrated with lysis buffer. The resulting complexes were analyzed for their dimerization content using mass photometry (below)

### Mass photometry

To prepare chambers, the microscope cover glass (#1.5 24 × 50 mm, Deckgläser) was cleaned by rinsing it with water, followed by isopropanol, and repeating this rinsing process three times. At the end, the slide was air dried by nitrogen. CultureWell silicone gaskets (Grace Bio-Labs) were cut and placed onto the freshly cleaned cover glass. Samples and standard proteins were diluted in lysis buffer (50 mM HEPES, 150 mM KCl, 1 mM MgCl2, 3 mM β-mercaptoethanol, 5% glycerol). For calibration, standard proteins BSA (Sigma), Apoferritin (Sigma), and Thyroglobulin (Sigma) were diluted to 10–50 nM. The no-specific binding events of single molecules on cover glasses were scattered, and the masses of these molecules were measured by the OneMP instrument (Refeyn) at room temperature. Data were collected at an acquisition rate of 1 kHz for 100 s by AcquireMP (Refeyn) and subsequently analyzed by DiscoverMP (Refeyn). For each concentration of recombinant proteins, the measurement was performed three times and repeated with two different protein preparations. Results from one representative measurement are shown in figures.

### Cryo-EM sample preparation and imaging

SEC purified complexes of TBC-DEG-β-tubulin and TBC-DE-β-tubulin were concentrated to 0.6 mg/mL. Concentrated TBC-DEG-β-tubulin was also incubated with bispecific crosslinker with 200 nM BS3 (bis-sulfosuccinimidyl-suberate) (Thermofisher) for two hours to increase the abundance of the back-to-back dimer state. Copper Quantifoil R1.2/1.3 grids were glow-discharged, and 4μl of sample buffer exchanged using a dialysis cassette into 50 mM HEPES, 150 mM KCl, and 0.01% NP40 was applied to glow discharged grids inside the chamber of a Mark III Vitrobot (ThermoFisher) at 20°C and 100% humidity. The grid was incubated with sample for 30 seconds. Sample was then blotted at force 8 for 3-5 seconds and then plunged into liquid ethane. TBC-DE-β-tubulin grids were screened, and data was collected using a ThermoFisher Glacios operated using a Gatan K3 direct electron detector collecting 60 frames per 2 sec for a total dose of 80 e-/Å. Data was collected using super resolution mode at 0.44Å/pix. Two TBC-DEG-β-tubulin grids were imaged in differing conditions. The first was using a Thermofisher Krios microscope operated using a Gatan K3 direct electron detector collecting 40 ms frames at 1.6-2.0 s exposure time for a total dose of 50 e-/Å. Data was collected in counting mode at 0.83Å/pix. The second was using a Thermofisher Krios operated using a Gatan K3 direct electron detector 40 ms frames at 1.6-2.0 s exposure time for a total dose of 50 e-/Å. The data was collected in counting mode at 1.06 Å/pix.

### Single particle analysis pipeline

For each dataset of TBC-DEG-β-tubulin and TBC-DE-β-tubulin described in **Table 1**, image movies were motion corrected through RELION suite Motioncor2 using a 5X7 patch and B-factor of 150. Images were picked by LoG Picker, extracted, and were subjected to 2D classification using Cryosparc 4.1(*25*). Multiple rounds of 2D-Classification were used to remove junk particles. Upon 2D classification of both datasets revealed the presence of dimeric assemblies. For initial structures, 2D classes were visually separated and *ab initio* structures were generated in Cryosparc. After *ab initio*, particle image data was subjected to a heterogeneous refinement to further classify the data into monomers or dimers. 2D classification was repeated to ensure the fidelity of monomer and dimer particle pools. After heterogeneous refinement, each particle dataset was treated differently based on oligomerization state. For the pooled TBC-DE-β-tubulin monomer particles (**Figure S3A**), the most populated heterogeneous refined class was subjected to a homogenous refinement, non-uniform refinement, Local CTF refinement, and a local refinement. The pooled dimer classes of TBC-DE-β-tubulin were subjected to a heterogeneous refinement of their *ab initio* produced classes and a homogeneous refinement (**Figure S3A**).

For the dataset 1 of TBC-DEG-β-tubulin, the three *ab initio* classes were subjected to a heterogeneous refinement (**Figure S2A**). The best class was subjected to a homogeneous refinement, local CTF refinement, and a local Refinement. This best class contained the core structure of TBCD, Arl2, and β-tubulin (**Figure S2A**). There was a noisy region which was determined to be TBCE based on the positioning of the noisy region relative to the core when comparing it to the TBC-DE-β-tubulin structure (**Figure S2A**). The whole structure was masked and subjected to a 3D-classification with four classes, filtered to 7 angstroms, and with hard classification activated. Each class was subjected to a local refinement, and the best class was chosen for segmentation of the TBCE lever arm density (**Figure S2A**). That density was composited with the earlier obtained core structure of TBCD, Arl2, and β-tubulin. For dataset 2 of TBC-DEG-β-tubulin, the dimer particle pool was subjected to an *ab initio* job with three classes, and the resulting classes were subjected to a heterogeneous refinement. The best class was then further refined by a homogenous refinement, local CTF refinement, and a local refinement. The final refined map resolution maps and Fourier shell correlation (FSC) curves resulting from each dataset are presented (**Figure S2B-D; Figure S3B-D**)

### Model building and refinement

All human TBC-DEG-β-tubulin and TBC-DE-β-tubulin maps were modeled using a combination of ISOLDE, Coot(*26*) and PHENIX(*27*), starting with the AlphaFold3 models for human TBCD, TBCE, and Arl2 and the Trichoplusia ni β-tubulin(*22*). AlphaFold3 models were initially fit using ChimeraX and morphed using ISOLDE with secondary structure restraints. The models were subjected to cycles of Coot-based manual building of loops and side chain corrections and real space refinements in PHENIX. All TBCE-LRR and CapGly domains side chains were truncated, but residue assignments were kept due to the registry being evident by the well-defined ubiquitin domain. Dimer models were all composed of fit monomer models, but with side chains truncated due to medium resolution of density maps. The final model validation was performed in PHENIX (**Table 1**). Models and maps are deposited into the RSCB and EMDB (EMDB ID: XXXX, PDB ID: XXXX), and refinement and validation details are described in **Table 1**. Figures were generated using ChimeraX(*28*).

### AlphaFold3 model predictions

To determine TBC-DEG-β-tubulin AlphaFold3 models, sequences for the human TBCD, TBCE, Arl2, and Trichoplusia ni porcine β-tubulin and one GTP molecule were entered into a single multi-subunit determination using the AlphaFold3 server (www.alphafoldserver.com)(22). A single representative model is presented in figure XXX with pIDDT values per residue displayed and their corresponding PAE matrix with accuracy of residue position error. To determine TBC-DE-β-tubulin AlphaFold3 models, sequences for the human TBCD, TBCE, Trichoplusia ni β-tubulin, and one GTP molecule was entered into a single multi-subunit determination using the AlphaFold3 server (www.alphafoldserver.com)(22). All five AlphaFold3 models were comparable in PAE value. A single representative model is presented in figure XXX with the moderate to high confidence pIDDT values per residue displayed and their corresponding PAE matrix with accuracy of PAE per residue.

## Supporting information

https://www.dropbox.com/scl/fi/s9kgf2e5tfx14d8ef11m0/formated_supplementary_materials_revised.pdf?rlkey=d67chbew82lo49o3wptt5hewv&dl=0

## Acknowledgements

We acknowledge the support and advice of Dr Fei Guo and the BIOEM facility (Molecular Cellular Biology, UC-Davis) and for support in cryo-EM data collection. We acknowledge the extensive support for cryo-EM data collection from the National Center for CryoEM Access and Training (NCCAT) and the Simons Electron Microscopy Center located at the New York Structural Biology Center. NCCAT is supported by the NIH Common Fund Transformative High Resolution Cryo-Electron Microscopy program (U24 GM129539, and NIGMS R24 GM154192) and by grants from the Simons Foundation (SF349247) and NY State Assembly. We are grateful for the support and diligence of the NCCAT staff in providing expertise to identify best conditions for cryo-EM imaging. We thank Dr Camille Scott at the UC-Davis High Performance Computing facility for computational support. We thank Dr Bharti Singal (Stanford Cryo-EM center) for support on some initial cryo-EM data processing. We thank Dr Gant Luxton (Molecular Cellular Biology, UC-Davis) for the critical reading of this manuscript. JAB acknowledges grant support from the National Institutes of Health (GM110283, GM158334).

## Author Contribution statement

AT carried out biochemical studies, prepared cryo-EM grids, collected cryo-EM data, determined and refined all structures, built and refined all models, co-wrote and co-revised manuscript. MA prepared and screened cryo-EM grids. VG carried out biochemical studies. JAB conceived the project, provided grant support, supervised and advised all co-authors, carried out biochemical experiments, prepared figures, wrote and revised manuscript.

## Data Availability Statement

Cryo-EM maps and models presented here will be available in the Electron microscopy database (EMDB) with the EMBD-IDs: EMD-74878, EMD-74875, EMD-74462, EMD-74461, EMD-74460, EMD-74459, EMD-74458. The corresponding models are available at the Protein data bank (PDB) with the accession numbers 9ZNV,9ZNW,9ZNX,9ZNY,9ZNZ

## Competing interest statement

The authors declare no competing interests

## Notes

### Competing Interest Statement

The authors have declared no competing interest.

## References

1. A. Akhmanova, L. C. Kapitein, Mechanisms of microtubule organization in differentiated animal cells. Nat Rev Mol Cell Biol 23, 541–558 (2022).

2. A. Akhmanova, M. O. Steinmetz, Control of microtubule organization and dynamics: two ends in the limelight. Nat Rev Mol Cell Biol 16, 711–726 (2015).

3. N. B. Gudimchuk, J. R. McIntosh, Regulation of microtubule dynamics, mechanics and function through the growing tip. Nat Rev Mol Cell Biol 22, 777–795 (2021).

4. M. Hopfler et al., Mechanism of ribosome-associated mRNA degradation during tubulin autoregulation. Mol Cell 83, 2290–2302 e2213 (2023).

5. Z. Lin et al., TTC5 mediates autoregulation of tubulin via mRNA degradation. Science 367, 100–104 (2020).

6. D. Gestaut et al., Structural visualization of the tubulin folding pathway directed by human chaperonin TRiC/CCT. Cell 185, 4770–4787 e4720 (2022).

7. S. A. Lewis, G. Tian, N. J. Cowan, The alpha- and beta-tubulin folding pathways. Trends Cell Biol 7, 479–484 (1997).

8. G. Tian, A. Bhamidipati, N. J. Cowan, S. A. Lewis, Tubulin folding cofactors as GTPase-activating proteins. GTP hydrolysis and the assembly of the alpha/beta-tubulin heterodimer. J Biol Chem 274, 24054–24058 (1999).

9. J. Al-Bassam, Revisiting the tubulin cofactors and Arl2 in the regulation of soluble alphabeta-tubulin pools and their effect on microtubule dynamics. Mol Biol Cell 28, 359–363 (2017).

10. S. Nithianantham et al., Tubulin cofactors and Arl2 are cage-like chaperones that regulate the soluble alphabeta-tubulin pool for microtubule dynamics. Elife 4, (2015).

11. D. A. Keays et al., Mutations in alpha-tubulin cause abnormal neuronal migration in mice and lissencephaly in humans. Cell 128, 45–57 (2007).

12. G. Tian et al., A pachygyria-causing alpha-tubulin mutation results in inefficient cycling with CCT and a deficient interaction with TBCB. Mol Biol Cell 19, 1152–1161 (2008).

13. G. Tian et al., Disease-associated mutations in TUBA1A result in a spectrum of defects in the tubulin folding and heterodimer assembly pathway. Hum Mol Genet 19, 3599–3613 (2010).

14. E. Flex et al., Biallelic Mutations in TBCD, Encoding the Tubulin Folding Cofactor D, Perturb Microtubule Dynamics and Cause Early-Onset Encephalopathy. Am J Hum Genet 99, 962–973 (2016).

15. N. Miyake et al., Biallelic TBCD Mutations Cause Early-Onset Neurodegenerative Encephalopathy. Am J Hum Genet 99, 950–961 (2016).

16. N. Martin et al., A missense mutation in Tbce causes progressive motor neuronopathy in mice. Nat Genet 32, 443–447 (2002).

17. R. Parvari et al., Mutation of TBCE causes hypoparathyroidism-retardation-dysmorphism and autosomal recessive Kenny-Caffey syndrome. Nat Genet 32, 448–452 (2002).

18. G. Tian, M. C. Huang, R. Parvari, G. A. Diaz, N. J. Cowan, Cryptic out-of-frame translational initiation of TBCE rescues tubulin formation in compound heterozygous HRD. Proc Natl Acad Sci U S A 103, 13491–13496 (2006).

19. X. B. Cai et al., Whole-exome sequencing identified ARL2 as a novel candidate gene for MRCS (microcornea, rod-cone dystrophy, cataract, and posterior staphyloma) syndrome. Clin Genet 96, 61–71 (2019).

20. A. Taheri, Z. Wang, B. Singal, F. Guo, J. Al-Bassam, Cryo-EM structures of the tubulin cofactors reveal the molecular basis of alpha/beta-tubulin biogenesis. Nat Commun (2025).

21. Y. Seong, H. Kim, K. Byun, Y. W. Park, S. H. Roh, Structural dissection of alphabeta-tubulin heterodimer assembly and disassembly by human tubulin-specific chaperones. Science 390, eady2708 (2025).

22. J. Abramson et al., Accurate structure prediction of biomolecular interactions with AlphaFold 3. Nature 630, 493–500 (2024).

23. M. J. Landrum et al., ClinVar: improving access to variant interpretations and supporting evidence. Nucleic Acids Res 46, D1062–D1067 (2018).

24. R. D. Vale, R. A. Milligan, The way things move: looking under the hood of molecular motor proteins. Science 288, 88–95 (2000).

25. A. Punjani, J. L. Rubinstein, D. J. Fleet, M. A. Brubaker, cryoSPARC: algorithms for rapid unsupervised cryo-EM structure determination. Nat Methods 14, 290–296 (2017).

26. A. Casanal, B. Lohkamp, P. Emsley, Current developments in Coot for macromolecular model building of Electron Cryo-microscopy and Crystallographic Data. Protein Sci 29, 1069–1078 (2020).

27. P. V. Afonine et al., New tools for the analysis and validation of cryo-EM maps and atomic models. Acta Crystallogr D Struct Biol 74, 814–840 (2018).

28. E. F. Pettersen et al., UCSF ChimeraX: Structure visualization for researchers, educators, and developers. Protein Sci 30, 70–82 (2021).

