## Supplementary material for "The Structural Basis of alpha/beta-tubulin Assembly and Disassembly by Tubulin Cofactors": https://www.dropbox.com/scl/fi/s9kgf2e5tfx14d8ef11m0/formated_supplementary_materials_revised.pdf?rlkey=d67chbew82lo49o3wptt5hewv&dl=0

### Figure S1-S14

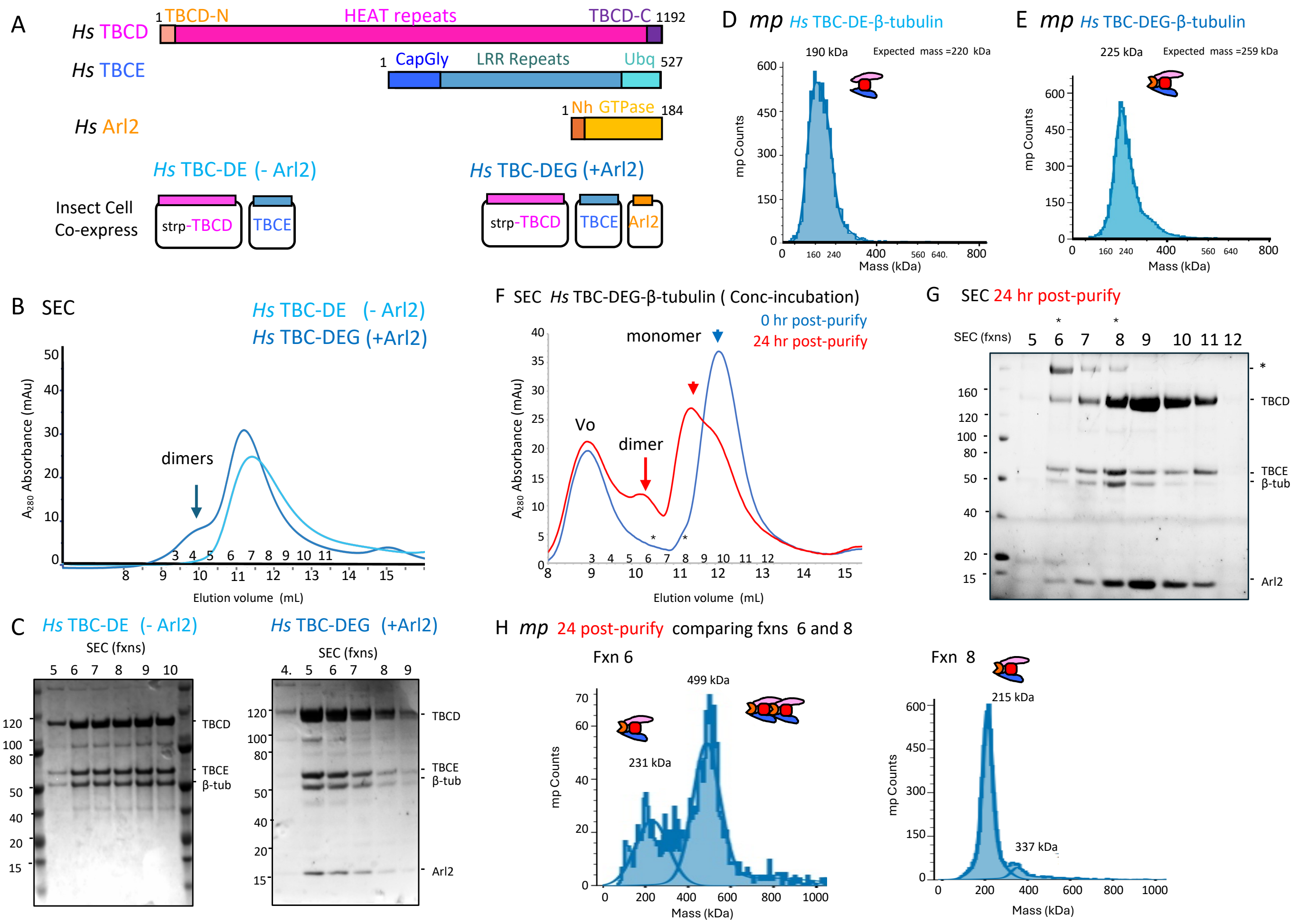


**Figure S1. Biochemical reconstitution of TBC-DE–β-tubulin and TBC-DEG–β-tubulin complexes purified from eukaryotic cells**

**A)** Domain organization of TBCD, TBCE, and Arl2.

**B)** Size-exclusion chromatography of freshly purified human TBC-DEG–β-tubulin (blue) and human TBC-DE–β-tubulin (cyan), with elution fractions indicated.

**C)** SDS–PAGE analysis of SEC fractions corresponding to chromatograms in panel B.

**D)** Mass photometry of freshly purified human TBC-DEG–β-tubulin.

**E)** Mass photometry of freshly purified human TBC-DE–β-tubulin.

**F)** Size-exclusion chromatography of concentrated human TBC-DEG–β-tubulin analyzed 1 hour (blue) or 24 hours (red) after incubation at 4 °C. Arrowheads mark monomeric peaks matching freshly purified samples in panel B; arrows mark dimeric species.

**G)** SDS–PAGE analysis of SEC fractions of TBC-DEG–β-tubulin after 24-hour incubation (red trace from panel F). The asterisk marks a non-dissociated complex containing TBCD, β-tubulin, and TBCE.

**H)** Mass photometry of fractions #6, #7, and #8 from panels F–G.
Left: fraction #6; middle: fraction #7; right: fraction #8.

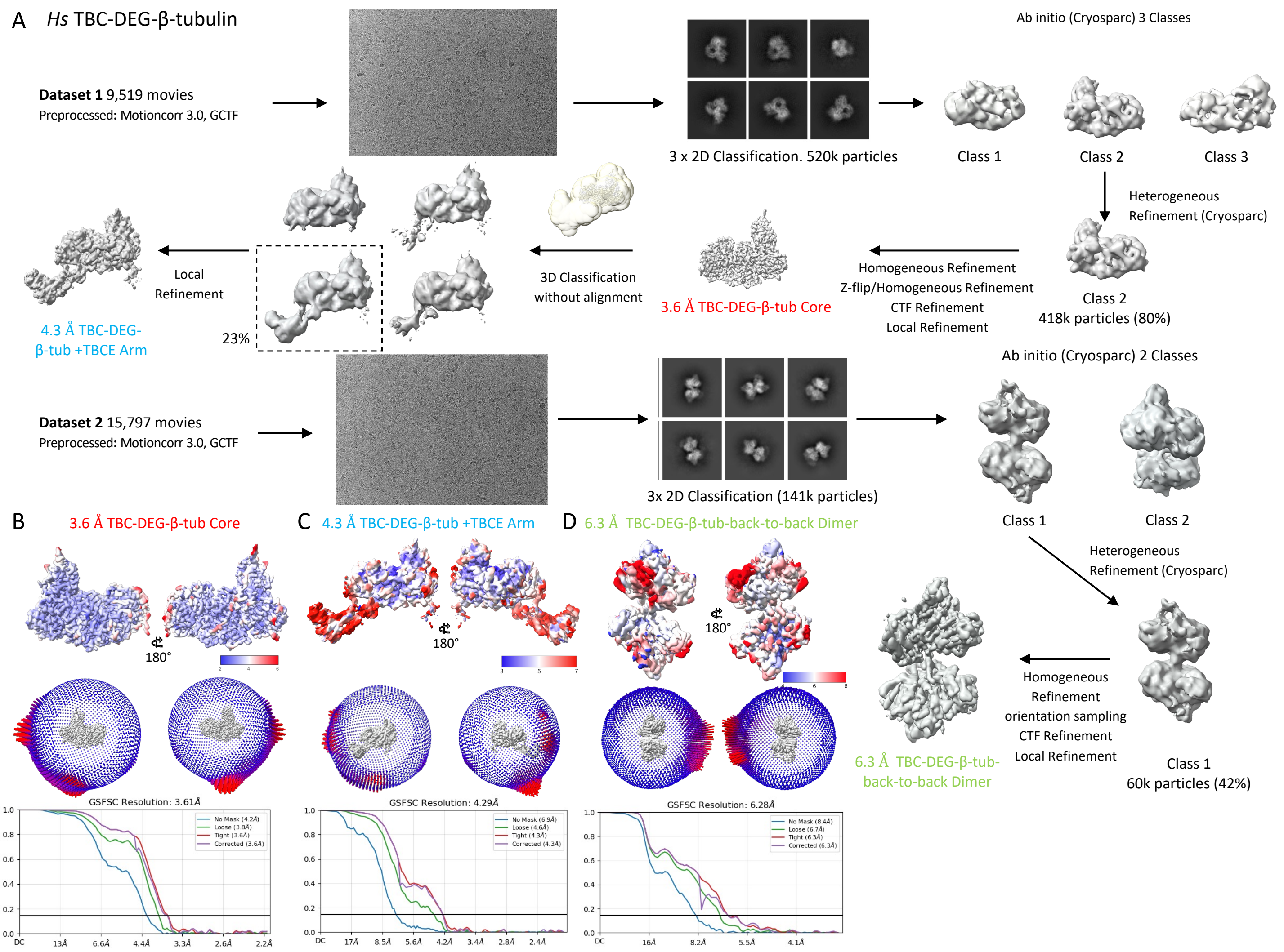


**Figure S2. Cryo-EM structure determination workflow for TBC-DEG–β-tubulin assemblies**

A) Overview of cryo-EM processing of Dataset 1 and Dataset 2 as described in materials and methods.

B) Top panel: Resolution-colored map (ResMap) for the 3.6 Å core. Middle panel: Angular distribution of particle views (two orientations). Bottom panel: Fourier Shell correlation (FSC) curve.

C) Top panel: Resolution-colored map of the 4.3 Å full TBC-DEG–β-tubulin structure. Middle panel: Angular distribution of views. Bottom panel: FSC curve.

D) Top panel: Resolution-colored map of the 6.3 Å back-to-back TBC-DEG–β-tubulin dimer map. Middle panel: Angular distribution of views. Bottom panel: FSC curve.

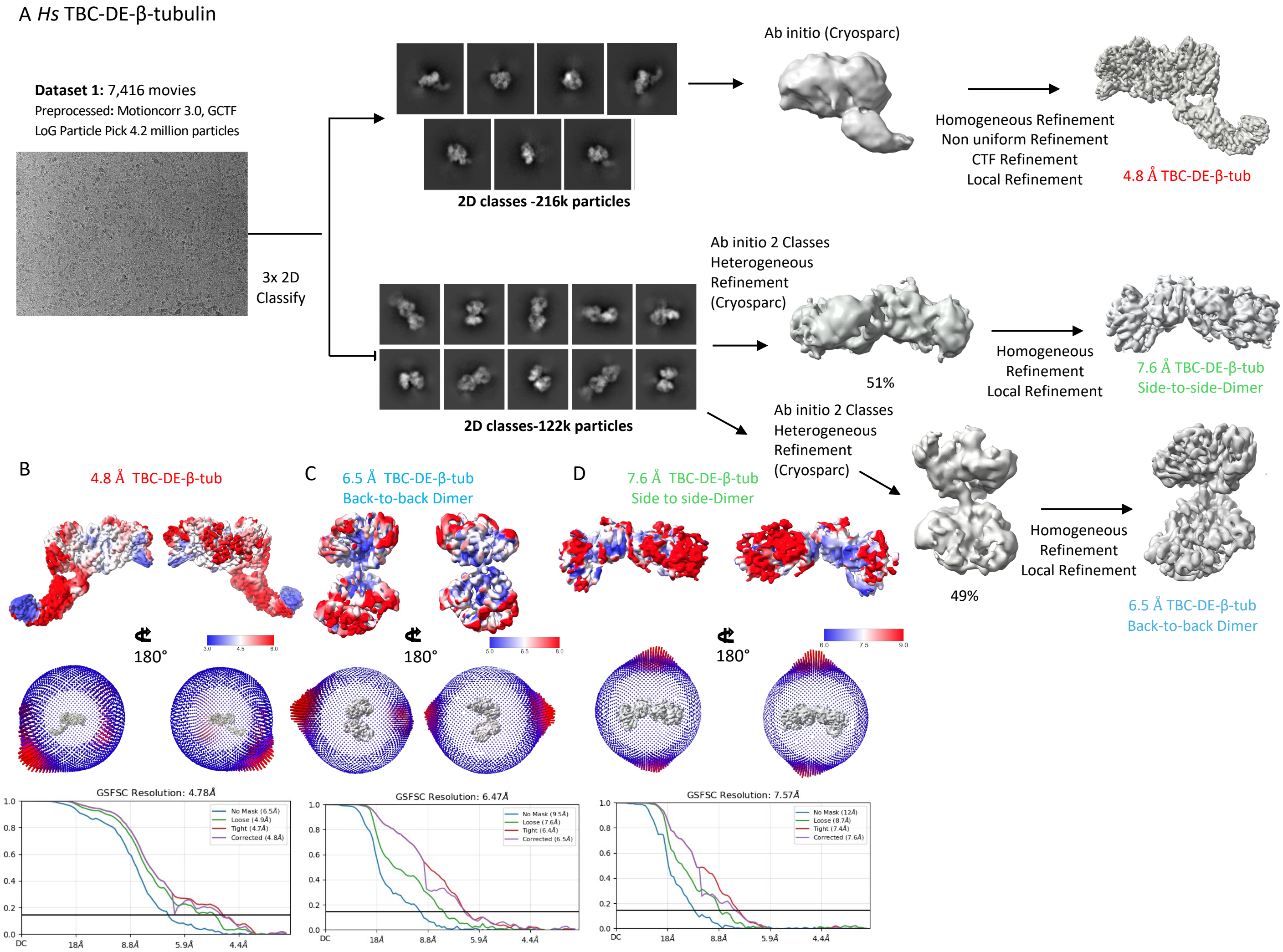


**Figure S3. Cryo-EM structure determination workflow for TBC-DE–β-tubulin assemblies**

A) Overview of Dataset 1 data processing for TBC-DE–β-tubulin as described in materials and methods.

B) Top panel: Resolution-colored map of the 4.8 Å TBC-DE–β-tubulin monomer. Middle panel: Angular distribution of particle views. Bottom panel: FSC curve.

C) Top panel: Resolution-colored map of the 6.3 Å back-to-back TBC-DE–β-tubulin dimer. Middle panel: Angular distribution of views. Bottom panel: FSC curve.

D) Top panel: Resolution-colored map of the 7.6 Å side-to-side TBC-DE–β-tubulin dimer. Middle panel: Angular distribution of views. Bottom Panel: FSC curve.

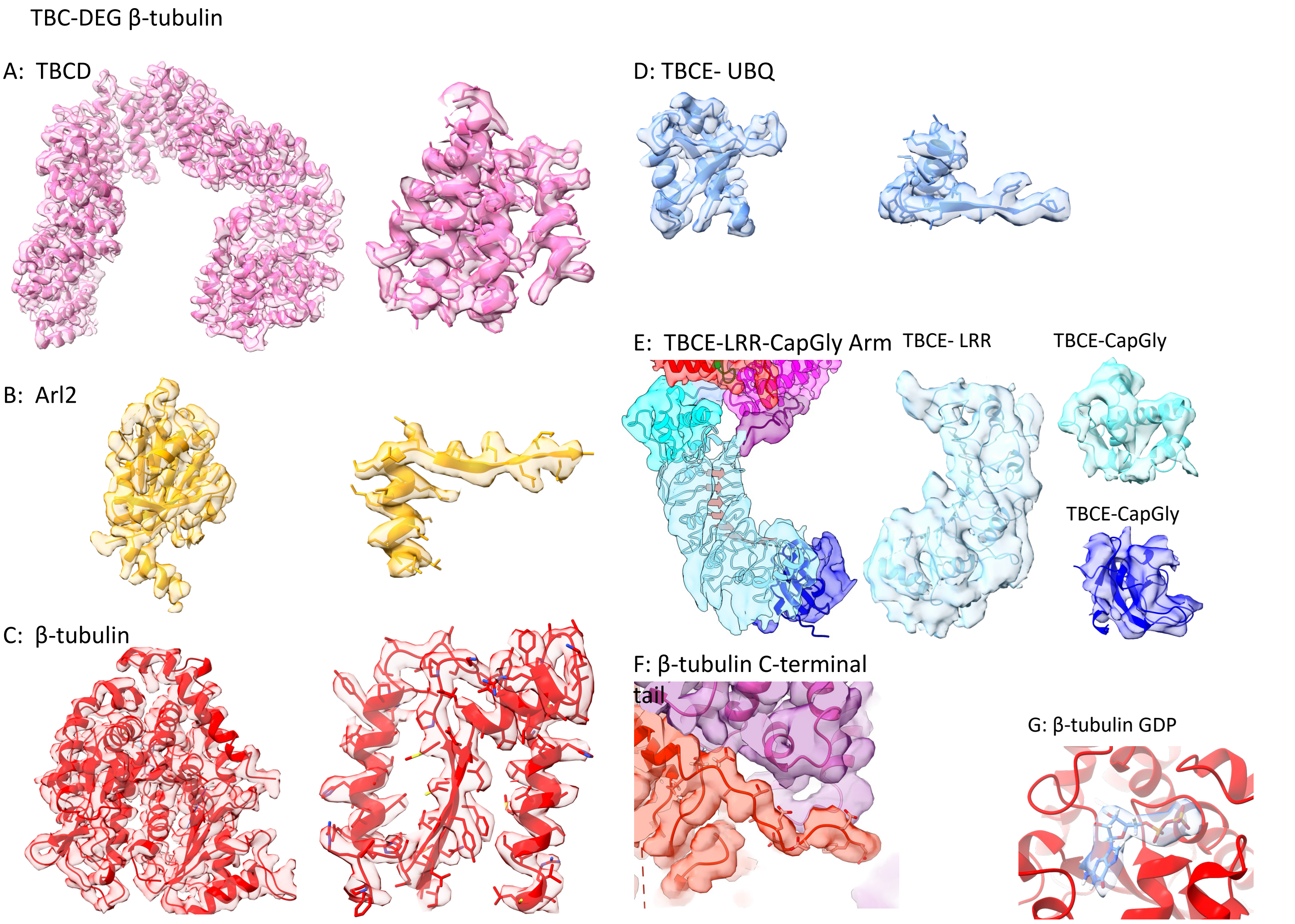


**Figure S4. Model quality and map-to-model fit for TBC-DEG–β-tubulin subunits and interfaces**

A) Left, full TBCD segmented density map and the model (pink). Right, close-up view on side chains
B) Left full Arl2 segmented density map and the model (orange). Right, close-up view on side chains.
C) Left full β-tubulin segmented density map and the model (red). Right, close-up view on side chains.
D) Left, TBCE-Ubq segmented density map and the model (cornflower). Right, close-up view on side chains.
E) Left, modeled density map for TBCE-LRR-CapGly arm. Right views of TBCE-LRR (sky blue), CapGly (blue) and 3HB (cyan) density map and models for these domains.
F) A close-up view of the β-tubulin C-terminal tail region (red) while bound to TBCD (pink)
G) A close-up view of the GDP (blue model) bound to β-tubulin (red).

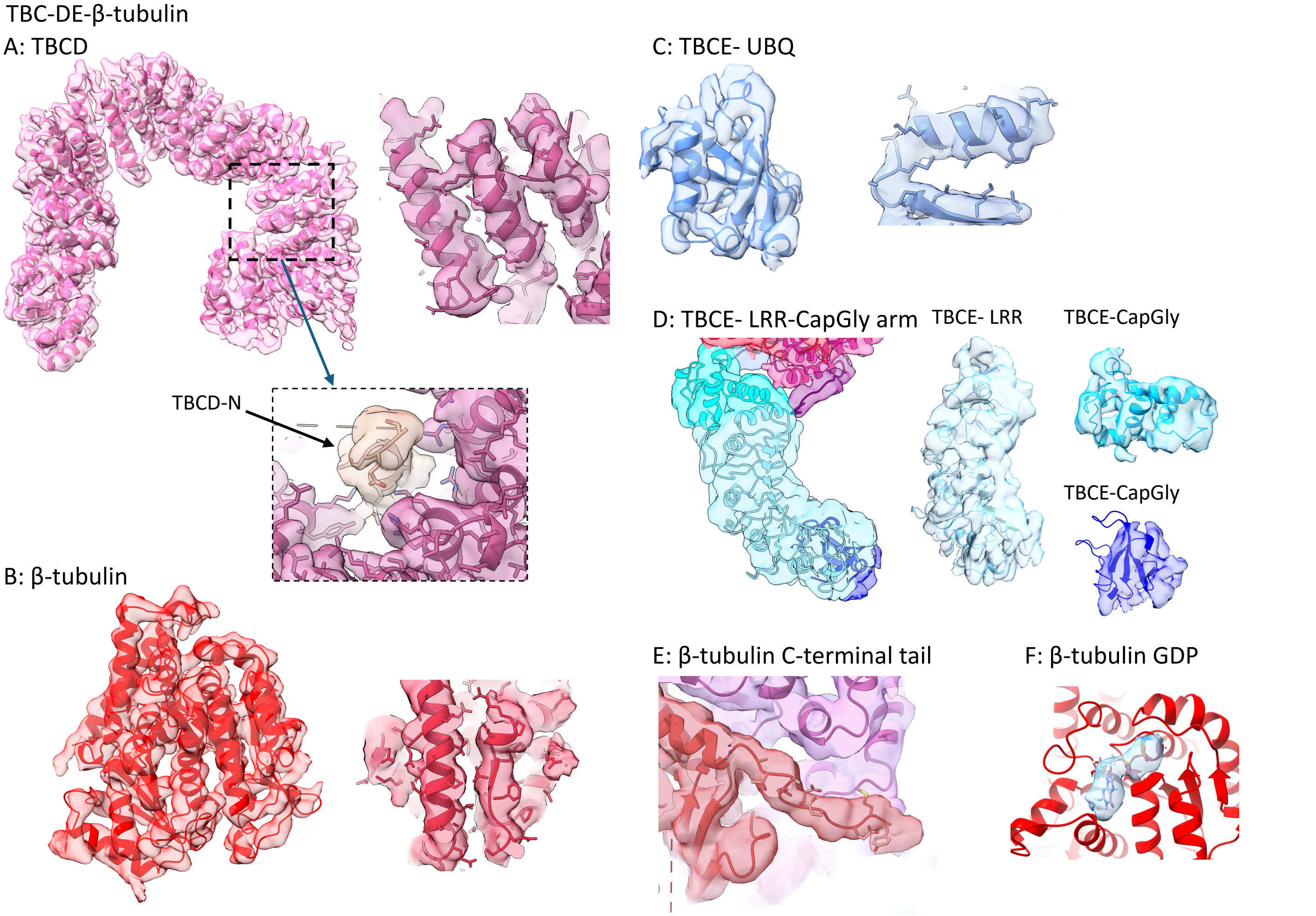


**Figure S5. Model quality and map-to-model fit for TBC-DE–β-tubulin subunits and interfaces**

A) Left, full TBCD segmented density map and the model (pink). Right, close-up view on side chains
B) Left full β-tubulin segmented density map and the model (red). Right, close-up view on side chains.
C) Left, TBCE-Ubq segmented density map and the model (cornflower). Right, close-up view on side chains.
D) Left, 5 Å map for TBCE-LRR-CapGly arm. Right views of TBCE-LRR (sky blue), CapGly (blue) and 3HB (cyan) density map and models for these domains.
E) A close-up view of the β-tubulin C-terminal tail region (red) while bound to TBCD (pink)
F) A close-up view of the GDP (blue model) bound to β-tubulin (red).

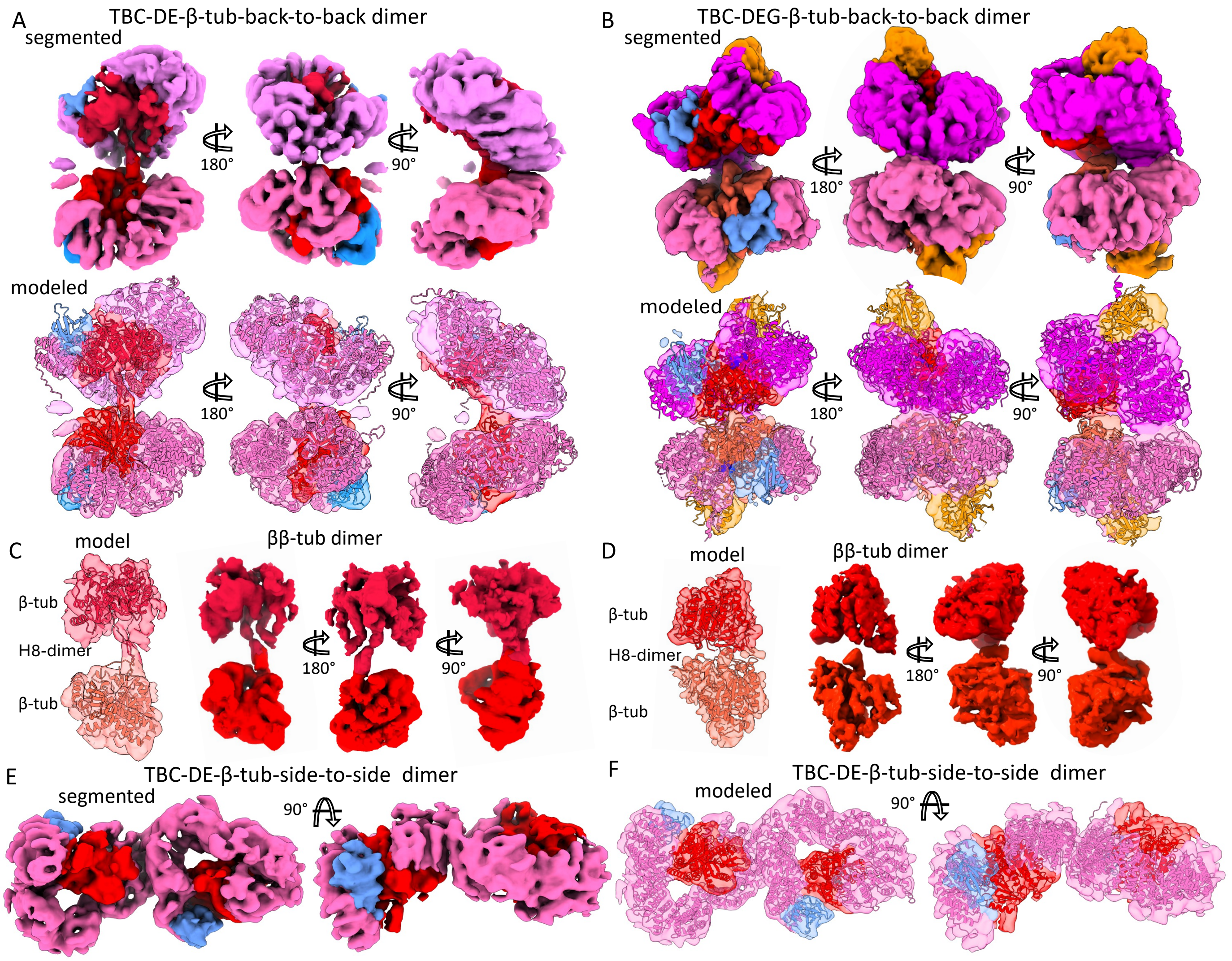


**Figure S6. Structural organization of TBC-DEG–β-tubulin and TBC-DE–β-tubulin back-to-back dimers and β–β-tubulin homodimer interfaces**

- **A)** Top panel, three rotated views for the segmented TBC-DE-β-tubulin back-to-back dimer with TBCD (pink), β-tubulin (red) and TBCE-ubq (cornflower blue). Bottom panel, three rotated views for the modeled density map for TBC-DE-β-tubulin colored as described above.
- **B)** Top panel, three rotated views for the segmented TBC-DEG-β-tubulin back-to-back dimer with TBCD (dark pink), Arl2 (orange), β-tubulin (red) and TBCE-ubq (cornflower blue). Bottom panel, three rotated views for the modeled density map for TBC-DEG-β-tubulin colored as described above.
- **C)** Left, modeled map of β-β-tubulin homodimer bound at the inside of TBC-DE-β-tubulin back-to-back dimer. The model shows each tubulin protomer shown in two shades of red. Right, three views of the segmented density of the β-β-tubulin homodimers.
- **D)** Left, modeled map of β-β-tubulin homodimer bound at the inside of TBCG-DE-β-tubulin back-to-back dimer. The model shows each tubulin protomer shown in two shades of red. Right, three views of the segmented density of the β-β-tubulin homodimers.
- **E)** Two rotated views for the segmented TBC-DE-β-tubulin side-to-side dimer with TBCD (pink), β-tubulin (red) and TBCE-ubq (cornflower blue).
- **F)** Two rotated views for the modeled density map for TBC-DE-β-tubulin side-to-side dimers, colored as described above in E.

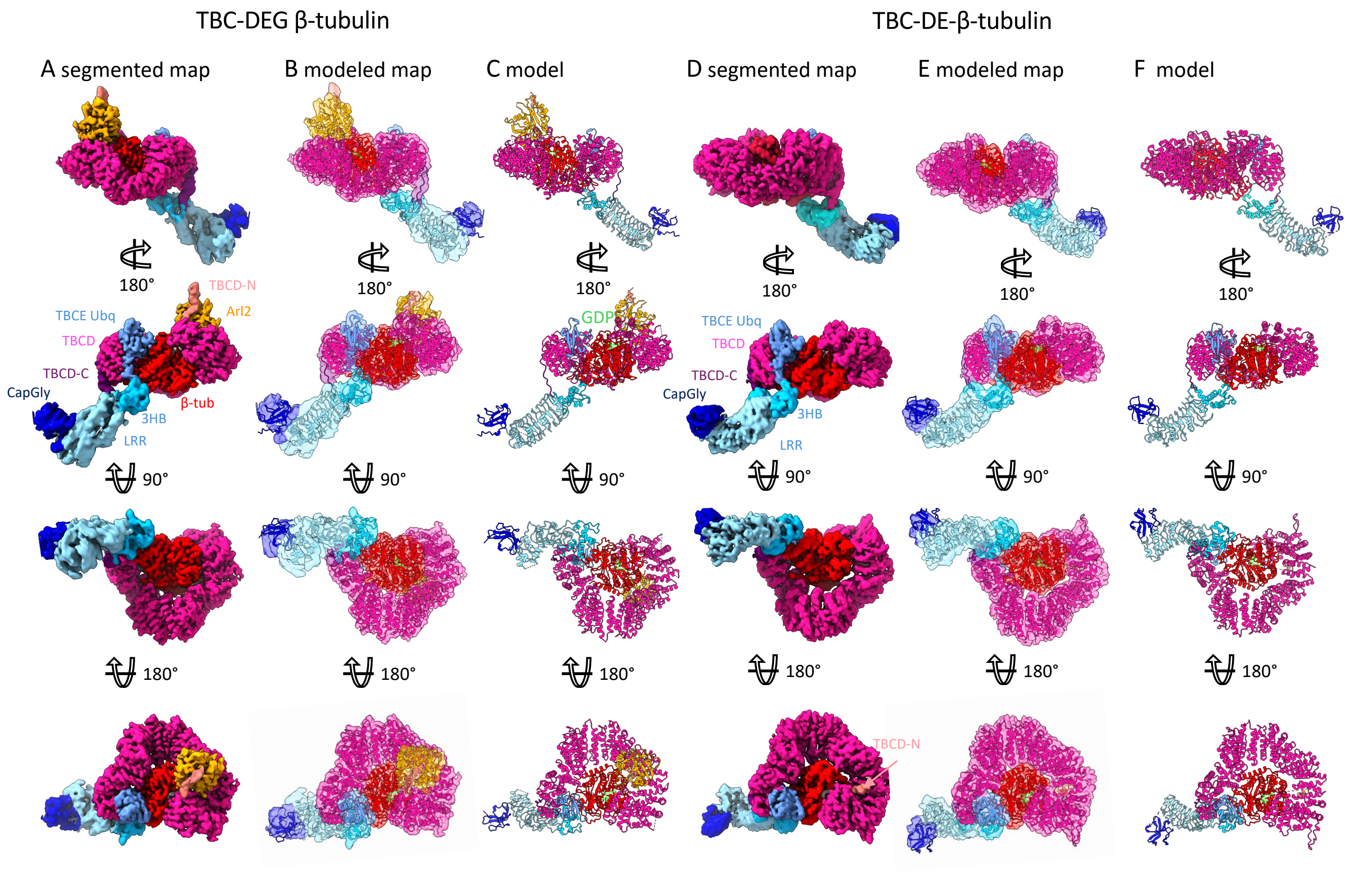


**Figure S7. Cryo-EM maps, models, and map-to-model correlation for TBC-DEG–β-tubulin and TBC-DE–β-tubulin structures**

A) Four rotated views (top to bottom) of the segmented TBC-DEG-β-tubulin map.
B) Four rotated views (top to bottom) of the modeled segmented TBC-DEG-β-tubulin map
C) Four rotated views (top to bottom) of the TBC-DEG-β-tubulin model
D) Four rotated views (top to bottom) of the segmented TBC-DE-β-tubulin map.
E) Four rotated views (top to bottom) of the modeled segmented TBC-DE-β-tubulin map
F) Four rotated views (top to bottom) of the TBC-DE-β-tubulin model

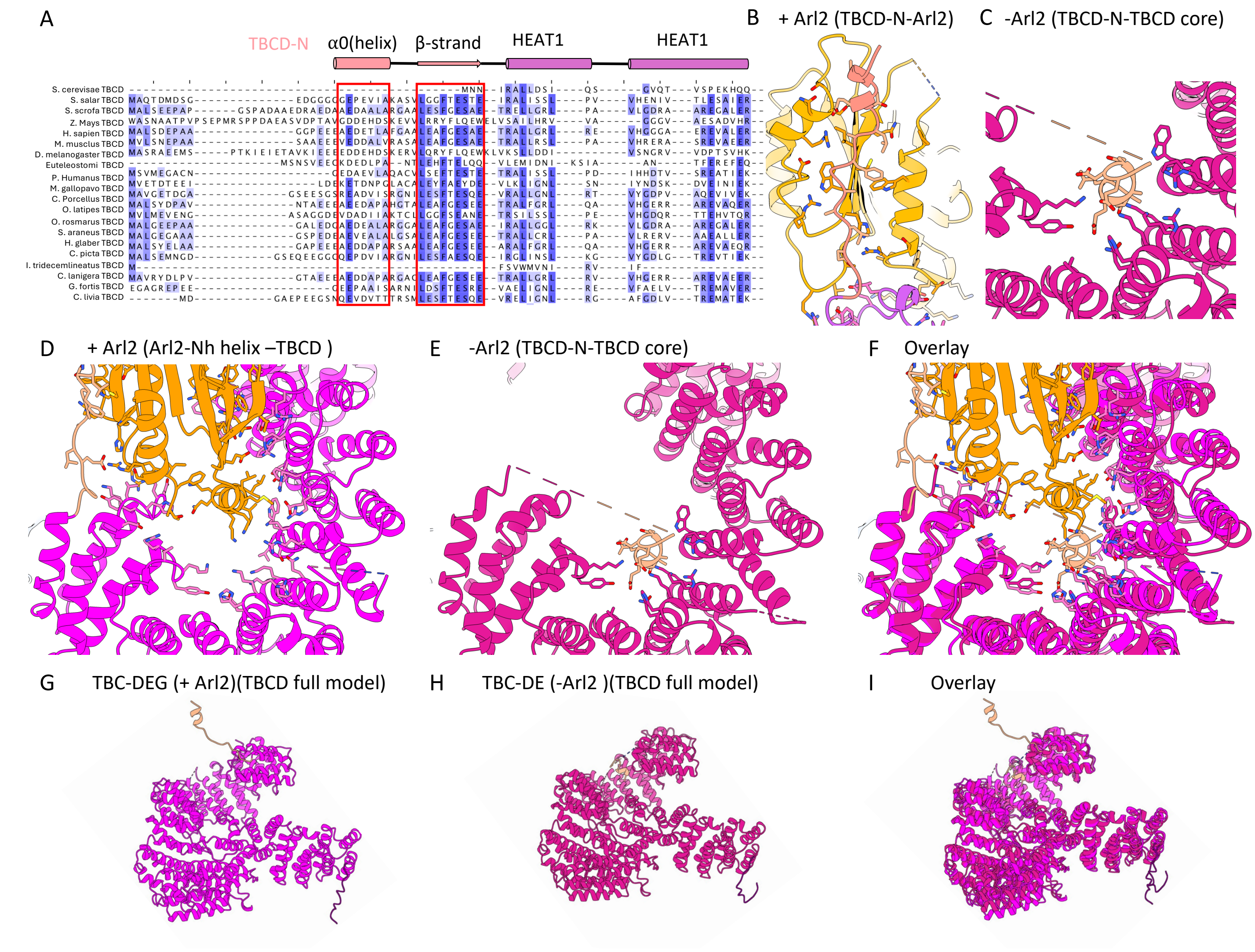


**Figure S8. Comparison of TBCD in TBC-DE–β-tubulin and TBC-DEG–β-tubulin structures and the role of the TBCD-N/Arl2 interface**

A) **S**equence conservation of the TBCD extreme N-terminus (TBCD-N) domain reveals the conservation of the α0-helix and the strand prior to HEAT1. Sequences for α0 and β-strand marked by red boxes. Note the conservation of acidic residues in α0 and hydrophobic residues in β-strand.

B) Binding interactions of the TBCD-N helix in the Arl2-bound state in TBC-DEG-β-tubulin revealing ionic interactions between TBCD-N α0 and top of Arl2 GTPase and hydrophobic interactions between β-strand and Arl2 Central β-sheet using hydrophobic interactions.

C) Binding interactions of the TBCD-N helix to the TBCD turret in the TBC-DE-β-tubulin revealing ionic interactions between TBCD-N α0 helix and the TBCD-turret basic residues.

D) Top view of the residue interactions of Arl2 N-terminal helix (Nh) binding the TBCD turret.

E) Top view of the residue interactions of the TBCD-N α0 helix binding in the TBCD turret.

F) Overlay of the TBCD-N α-helix and Arl2 Nh helix binding at nearby sites within the TBCD turret.

G) Model for TBCD from the Arl2-bound TBC-DEG-β-tubulin structure

H) Model for TBCD from the TBC-DE-β-tubulin structure.

I) Overlay of TBCD models in both G and H.

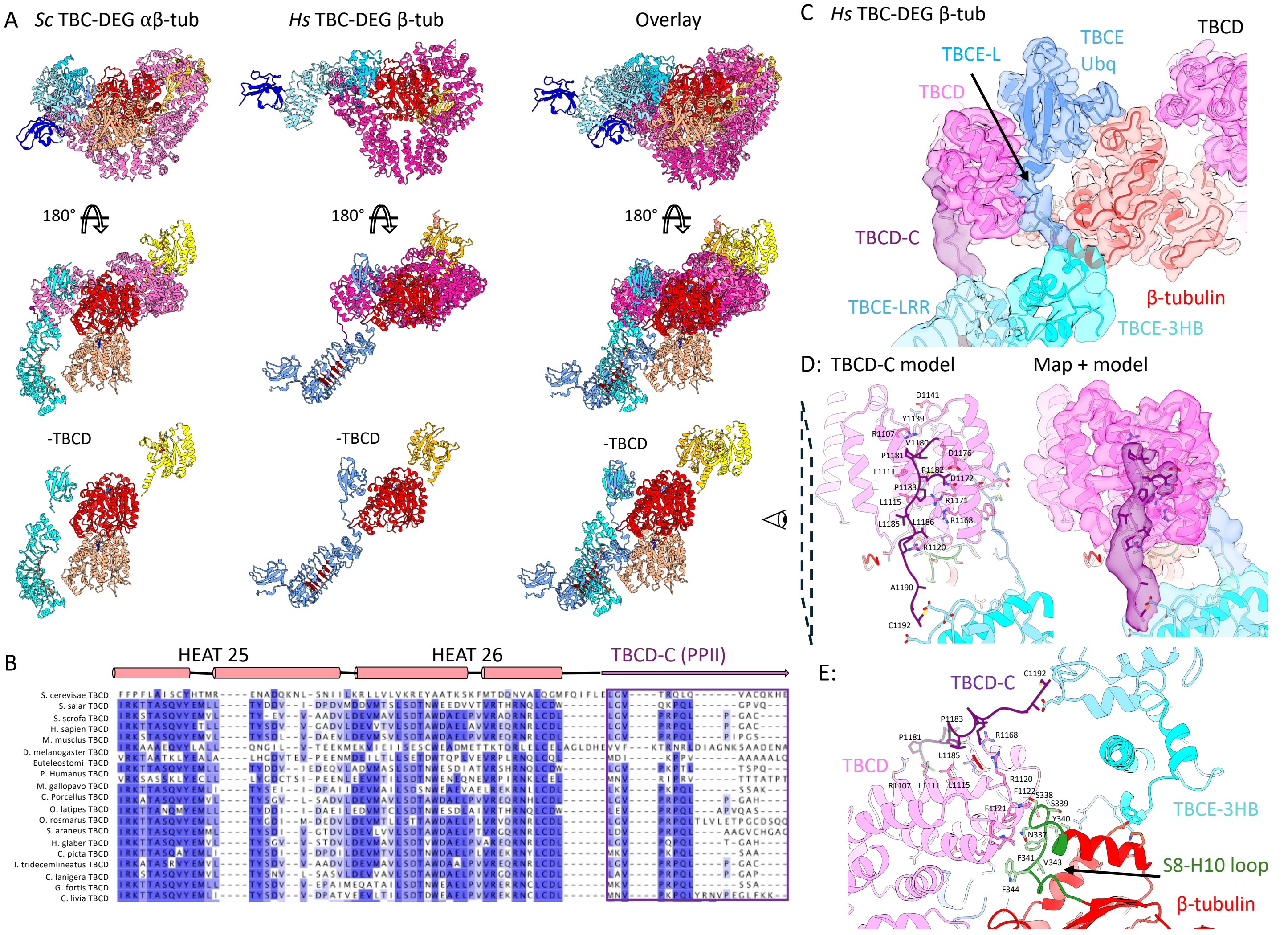


**Figure S9. Structural comparison of TBCE LRR-CapGly arm conformations in TBC-DEG–β-tubulin versus TBC-DEG–αβ-tubulin**

A) Top panels, Top view comparison of the yeast TBC-DEG–αβ-tubulin (left) and human TBC-DEG–β-tubulin (middle) and overlay of both (left) highlighting the Arl2 and TBCE conformational changes. Second panels, 90° rotated view of the top panels. Bottom panel, models shown in the middle panels with the TBCD models removed.

B) Sequence conservation analysis of the TBCD C-terminal region and including the TBCD HEAT26 and TBCD-C poly proline II (PPII) region.

C) Segmented Map and model fit of the TBCD-TBCE-β-tubulin interface site including The TBCD C-terminal HEAT repeats, TBCD-C, TBCE Ubq domain, TBCE-L linker region along the TBCD-C-terminus and TBCE-3HB.

D) Left, Model view of the polyproline-rich C-terminus of TBCD and its interaction with TBCE. Right, map and model view for the panel on the right.

E) Bottom model view of the β-tubulin H10 S8 loop rearrangements stabilized by the TBCD C-terminus and its interface with the TBCE LLR and TBCD-C stabilization of the swung-out state.

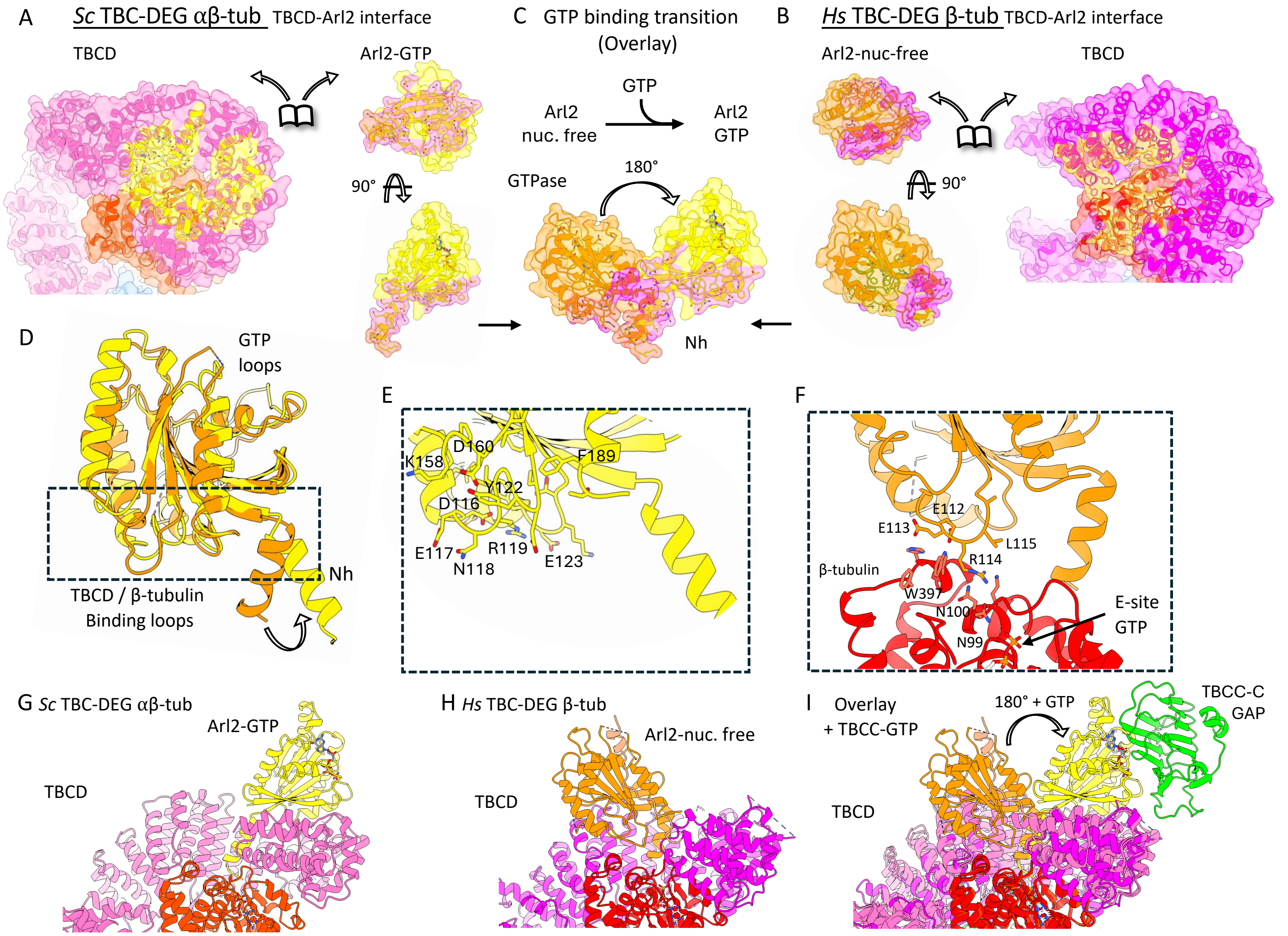


**Figure S10. Structural transitions in the Arl2 GTPase domain comparing TBC-DEG–β-tubulin and TBC-DEG–αβ-tubulin**

A) A dissociated surface view of the Arl2-binding interfaces in yeast TBC-DEG–αβ-tubulin. Left panel, view of the TBCD with Arl2 footprint marked in yellow. Top Right panel, shows a opposite view of Arl2 with TBCD footprint. Bottom right panel, 90° rotated showing a side view of Arl2.

B) A dissociated surface view of the Arl2-binding interfaces in yeast TBC-DEG–β-tubulin. right panel, view of the TBCD with Arl2 in nucleotide free w footprint marked in orange. Top left panel shows a opposite view of Arl2-nucleotide free state with TBCD footprint shown in pink. Bottom right panel, 90° rotated showing a side view of Arl2 nucleotide free.

C) Arl2 transitions in the GTP bound to nucleotide-free state comparing their states in A and. Conformational switch in Arl2 with and without nucleotide.

D) An overlay of the two Arl2 structures by their GTPase domain. Rearrangements of Arl2 Nh helix junction and the binding loops on TBCD and GTP-binding loops between the Nucleotide free (orange) and GTP state (yellow).

E) Close up view of the Arl2-GTPase in the yeast TBC-DEG–αβ-tubulin.

F) Close up view of the Arl2 GTPase interface with β-tubulin in human TBC-DEG–β-tubulin

G) Alternative view of Arl2-GTP state in the yeast TBC-DEG–αβ-tubulin.

H) Alternative view of the nucleotide-free Arl2 state in the human TBC-DEG–β-tubulin.

I) Alternative view of the TBCC-C interface with the Arl2 GTP state in relation to the Arl2- nucleotide state.

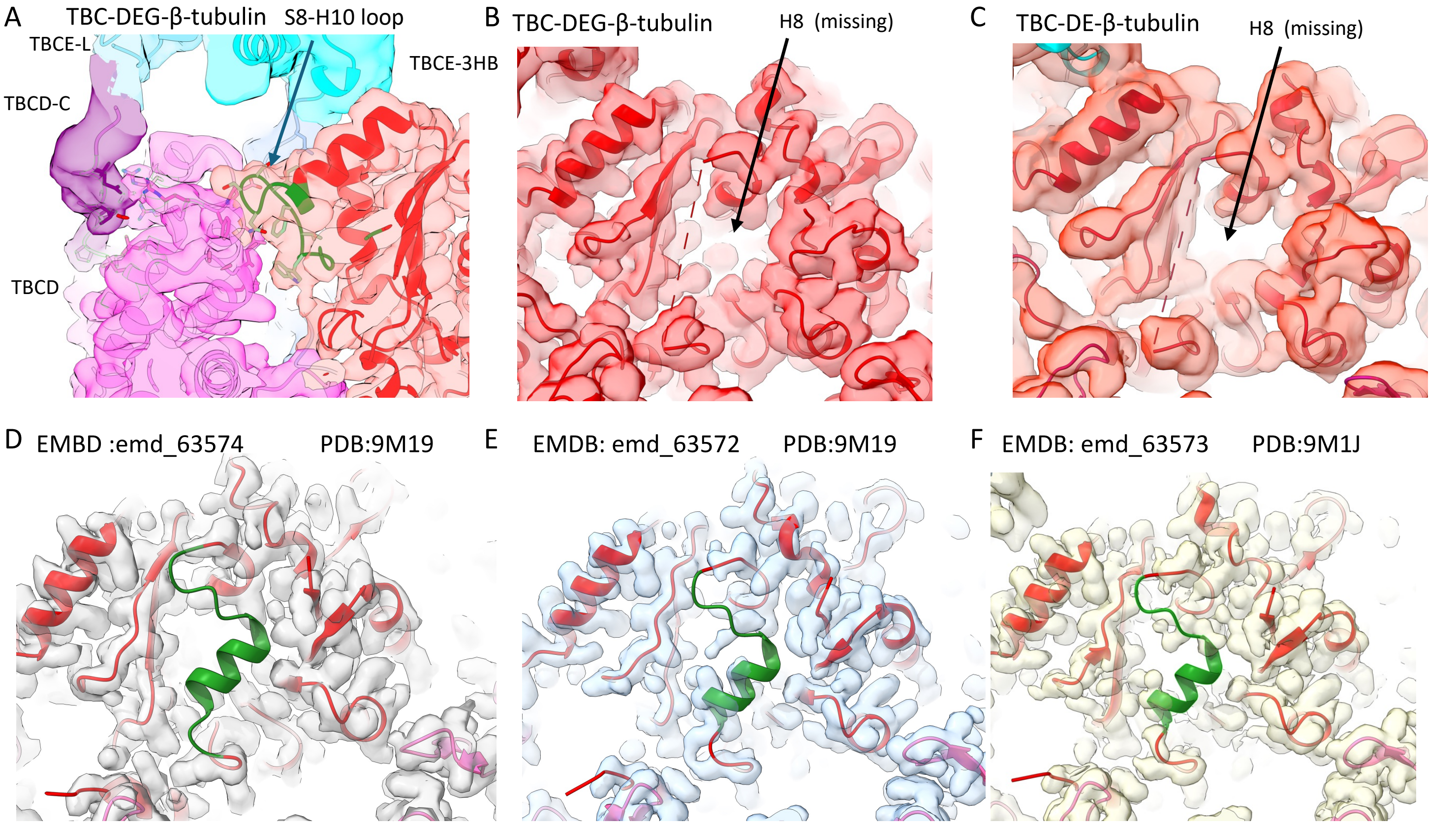


**Figure S11. Structural interfaces of the β-tubulin intradimer surface in TBC-DEG–β-tubulin and TBC-DE–β-tubulin structures**

A) A density map view of the rearrangement of the β-tubulin S8–H10 loop in TBC-DEG and TBC-DE structures.
B) A density view of the intradimer interface in β-tubulin revealing the absence of the H8 helix in TBC-DEG–β-tubulin.
C) A density view of the intradimer interface in β-tubulin revealing the absence of the H8 helix in TBC-DE–β-tubulin.
D) A density map view of the intradimer interface of deposited map EMDB 63574 and coordinates PDB 9M19 (*1*)
E) A density map view of the intradimer interface of deposited map EMDB 63573 and coordinates PDB 9M19 (*1*)
F) A density map view of the intradimer interface of deposited map EMDB 63572 and coordinates PDB 9M1J (*1*)

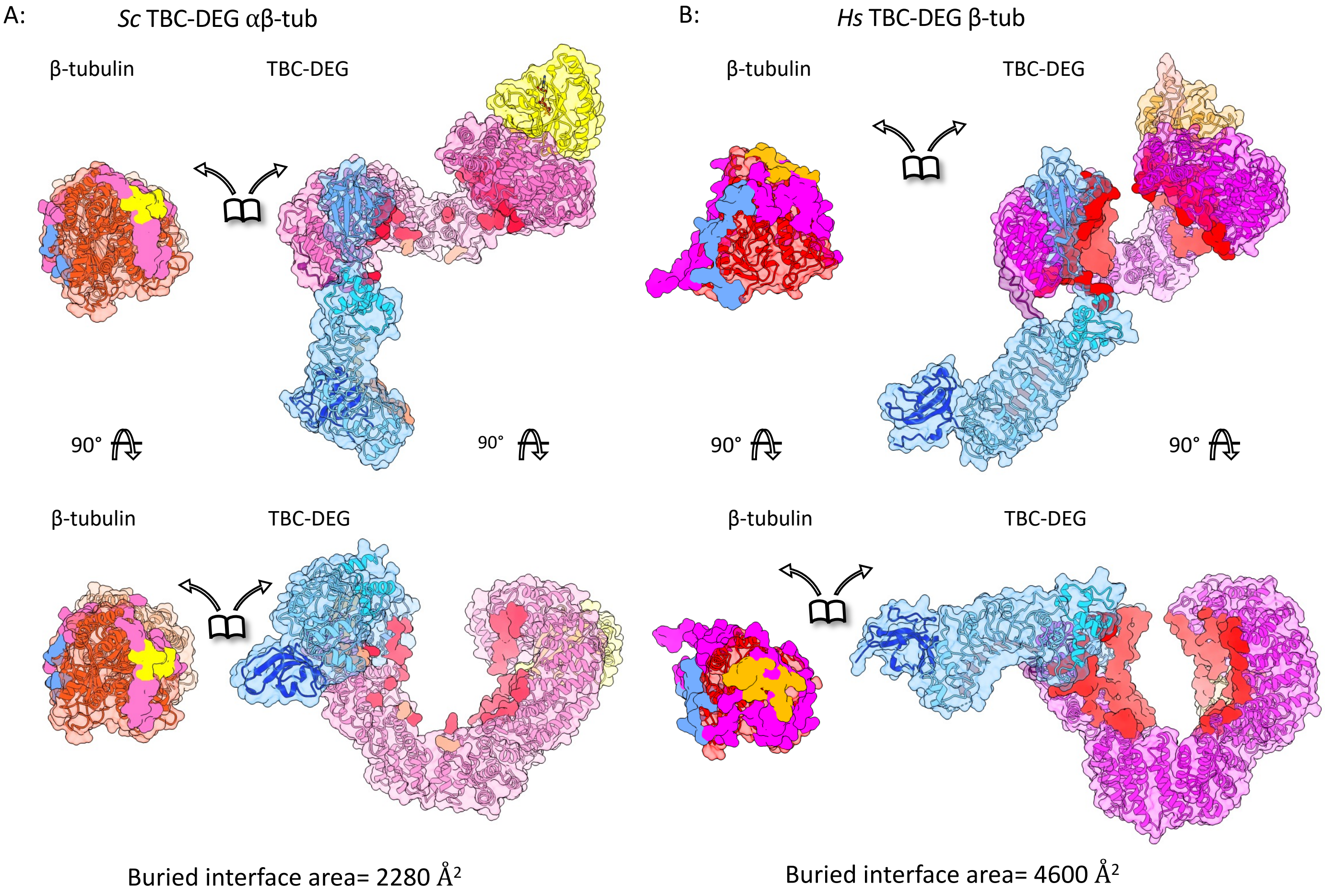


**Figure S12. Mapping of human disease mutations onto TBCD, TBCE, and Arl2 structures**

A) Two orthogonal views of the clinvar pathogenic mutations in TBCD, TBCE and Arl2 mapped onto the full TBC-DEG.

B) Top panel, Dissociated views of the TBCD, TBCE and Arl2 with the clinvar pathogenic mutations. Bottom, two top close-up views of TBCD and Arl2 showing the location of the clinvar mutations.

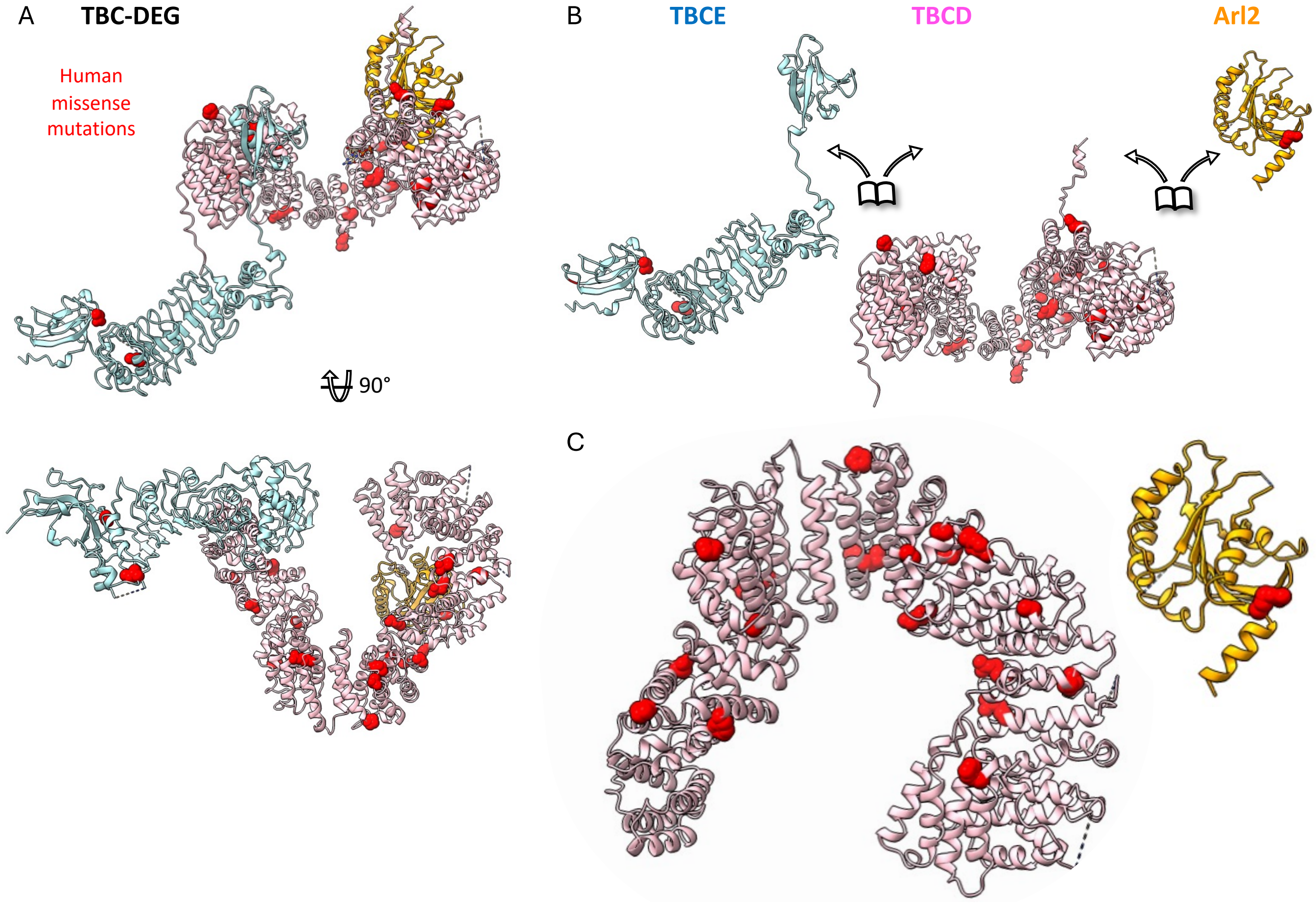


**Figure S13. Comparison of yeast and human TBC-DEG interaction interfaces with β-tubulin**

A) Two orthogonal views of the dissociated interaction interfaces of TBC-DEG of β-tubulin in the yeast TBC-DEG-αβ-tubulin structure. Top panels show top view and bottom panels show the side views.

B) Two orthogonal views of the dissociated interaction interfaces of TBC-DEG of β-tubulin in the yeast TBC-DEG-β-tubulin structure. Top panels show top view and bottom panels show the side views.

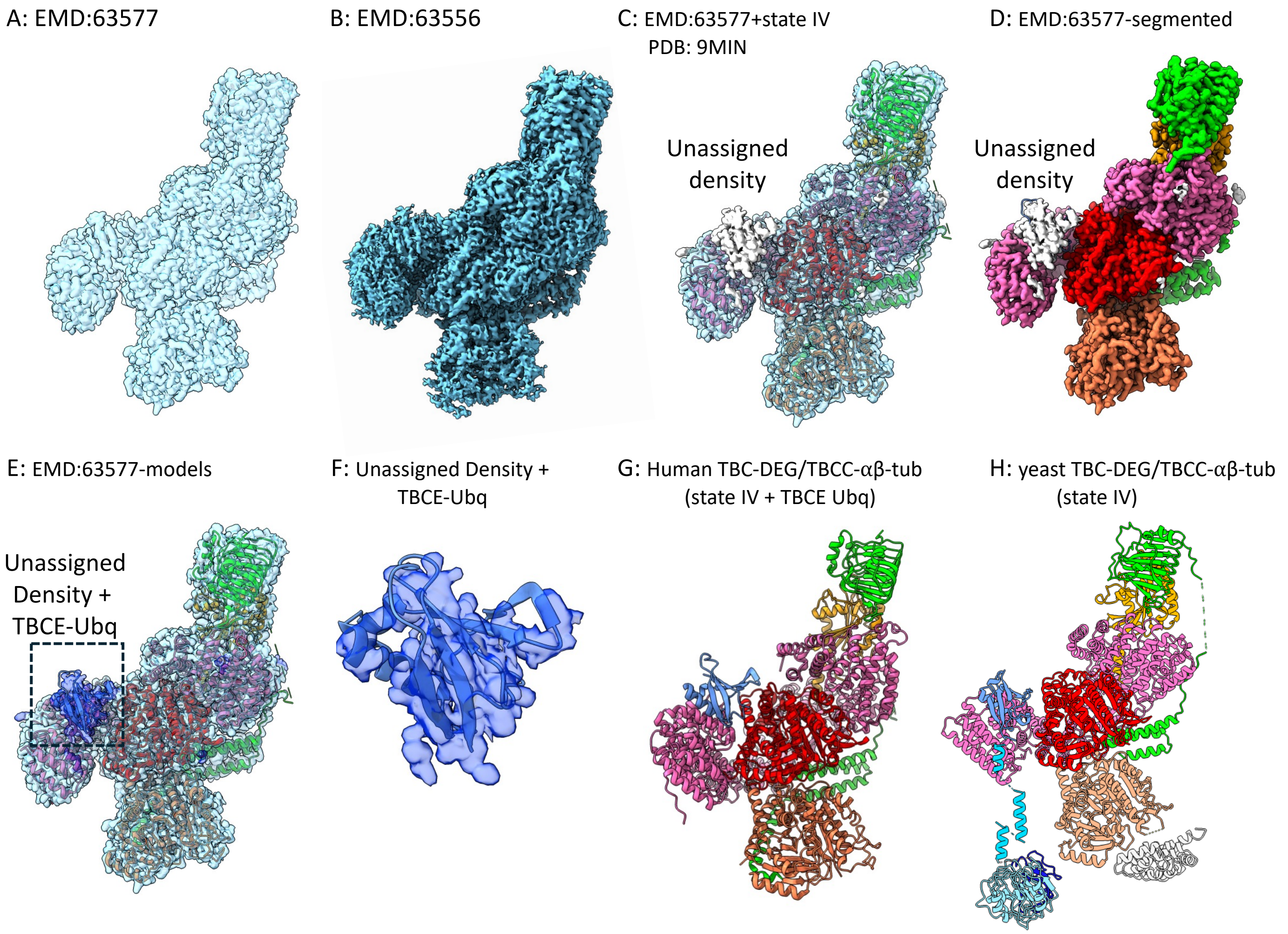


**Figure S14. Structural reanalysis of human TBC-DEG–TBCC–αβ-tubulin reveals unmodeled TBCE-Ubq domain density, supporting TBC-DEG stability throughout the catalytic cycle**

A) A side view of density map EMD:63577.
B) A side view of density map EMD:63556.
C) Segmented EMD:63577 highlighting unassigned density.
D) A fitted map view of segmented EMD:63577 fitted with PDB-ID 9MIN, highlighting unassigned density (white)
E) Segmented map view of EMD-63577 map with TBCE-Ubq domain fitted into added density (transparent blue)
F) Close-up of the TBCE-Ubq (blue) rigid-body fit into the additional unassigned density (blue)
G) View of the human TBC-DEG–TBCC–αβ-tubulin complex (state IV) showing the newly fitted TBCE-Ubq domain.
H) View of yeast TBC-DEG–TBCC–αβ-tubulin (state IV; EMD:63577) for comparison with panel G showing the full TBCE resolved (*2*).

**Supplementary References:**

1. Y. Seong, H. Kim, K. Byun, Y. W. Park, S. H. Roh, Structural dissection of alphabeta-tubulin heterodimer assembly and disassembly by human tubulin-specific chaperones. *Science* **390**, eady2708 (2025).
2. A. Taheri, Z. Wang, B. Singal, F. Guo, J. Al-Bassam, Cryo-EM structures of the tubulin cofactors reveal the molecular basis of alpha/beta-tubulin biogenesis. *Nat Commun*, (2025).
